# Structure-Function Dynamics of Engineered, Modular Neuronal Networks with Controllable Afferent-Efferent Connectivity

**DOI:** 10.1101/2022.11.24.517848

**Authors:** Nicolai Winter-Hjelm, Åste Brune Tomren, Pawel Sikorski, Axel Sandvig, Ioanna Sandvig

## Abstract

Microfluidic devices interfaced with microelectrode arrays have in recent years emerged as powerful platforms for studying and manipulating *in vitro* neuronal networks at the micro- and mesoscale. By segregating neuronal populations using microchannels only permissible to axons, neuronal networks can be designed to mimic the highly organized, modular topology of neuronal assemblies in the brain. However, little is known about how the underlying topological features of such engineered neuronal networks contribute to their functional profile. To start addressing this question, a key parameter is control of afferent or efferent connectivity within the network. In this study, we show that a microfluidic device featuring axon guiding channels with geometrical constraints inspired by a Tesla valve effectively promotes unidirectional axonal outgrowth between neuronal nodes, thereby enabling us to control afferent connectivity. Our results moreover indicate that these networks exhibit a more efficient network organization with higher modularity compared to single nodal controls. We verified this by applying designer viral tools to fluorescently label the neurons to visualize the structure of the networks, combined with extracellular electrophysiological recordings using embedded nanoporous microelectrodes to study the functional dynamics of these networks during maturation. We furthermore show that electrical stimulations of the networks induce signals selectively transmitted in a feedforward fashion between the neuronal populations. A key advantage with our microdevice is the ability to longitudinally study and manipulate both the structure and function of neuronal networks with high accuracy. This model system has the potential to provide novel insights into the development, topological organization, and neuroplasticity mechanisms of neuronal assemblies at the micro- and mesoscale in healthy and perturbed conditions.

## Introduction

Since the pioneering work by Taylor *et al*. in 2003, advancements in microfluidic technologies have been widely adopted by the neuroscience community for preclinical neuroscience research(1). The ability to segregate and integrate neuronal populations across compartmentalized microfluidic chips has facilitated the construction of robust, physiologically relevant model systems for studying neuronal networks *in vitro* with unparalleled experimental control. However, these platforms do not inherently aid the establishment of feedforward projection sequences. Such a topological organization is critical for promoting directional and hierarchical information processing, as can be seen in a range of brain areas, such as the laminar structure of the neocortex, the cerebrocerebellar loop and the entorhinal-hippocampal loop(2). Introducing physical constraints that accentuate the establishment of such network topologies could therefore trigger more diverse patterns of activity, input representations and adaptations in in *vitro* neuronal networks to better recapitulate the highly complex computational dynamics seen in vivo. While neuroengineering approaches have advanced significantly over the past two decades, there remains a need for more sophisticated model systems that efficiently promote directional axonal outgrowth, and as such the establishment of controllable afferent-efferent structural and functional connections.

*In vivo*, development of neuronal networks is tightly orchestrated by reciprocal, dynamic structure-function relationships shaped by an interplay between intrinsic neuronal self-organizing properties and spatiotemporally regulated chemical and physical guidance cues from the microenvironment(3–5). In *vitro*, dissociated neurons are seeded in a rather homogeneous environment lacking mechanical or chemical gradients to direct the establishment of directional projection sequences. As a solution to this, Dworak et *al*. pioneered the engineering of unidirectionally connected neuronal networks in 2010 by plating neurons sequentially in two chambers of a microfluidic device with a 10 days time difference(6, 7). This allowed axons from the presynaptic population to fill up the microchannels prior to the postsynaptic cells being plated. The study demonstrated that it is feasible to structure and study in *vitro* neuronal networks with controllable connectivity in microfluidic devices. However, while this approach proved efficient at promoting unidirectional axonal outgrowth, a considerable drawback with their approach is that neurons in the two populations will be at different maturation stages throughout an experiment. As *in vitro* neuronal networks gradually change activity patterns over time, such a developmental difference can have broad implications for the resulting network behaviour, and as such the physiological relevance of the model system(8). It also prevents investigations of complex cultures with more than two neuronal populations.

To allow cell populations to be plated concurrently, more recent studies have rather focused on manipulating the microenvironment of the cells to spatially control the growth dynamics of the neuronal assemblies. Over the past decade, several studies have demonstrated the effectiveness of microfluidic devices with geometrical constraints implemented within the microchannel architecture to steer the directionality of axonal outgrowth *in vitro*(9–17). Specific geometries embedded within the microchannels are used to promote axonal outgrowth in a single direction, while outgrowth in the opposite direction is impeded(18). Such topological constraints support the establishment of distinct pre- and post-synaptic populations of neurons, where axonal projections extend primarily in a single direction between the two populations. The first documented attempt at promoting unidirectional axon growth in this way was made by Peyrin *et al*. in 2011, demonstrating that narrow inlets at the postsynaptic side effectively reduced the chance of postsynaptic axons growing into the microchannels(9). This approach, however, significantly decreases the connectivity between the populations due to the narrow channel outlets.

An alternative approach has been to implement bottlenecks and traps inside the channels to coax the axons from the post-synaptic population away from the main channels and make them retract(10–13). A challenge with this approach is that it can cause axonal fragmentation and lead to debris build up inside the channels. To circumvent this, more recent studies have steered the afferent-efferent connectivity by implementing topographical features within the microchannels that guide the neurites from the presynaptic population through the channels, while the axons from the postsynaptic population are guided away from the main channels and back to their chamber of origin(14–16). While this approach shows great promise, examples of functional characterization of networks grown on such platforms are so far limited. While one of the latter studies demonstrated functional connectivity establishment between the populations using calcium imaging, the primary focus of all these three studies was on structural analysis of the networks using fluorescent indicators. While such analysis can give useful indications of the network structure, they do not necessarily capture all neuritic processes, and have inherent limitations in terms of quantitative analysis due to variations in fluorescence expression levels, and hence relevant intensity profiles(19). Integration of the microfluidic platforms with microelectrode arrays (MEAs) can allow for more robust analysis of how these microchannel architectures may influence network connectivity and activity(20).

To improve the robustness of engineered neuronal networks in *vitro* as models of neuronal networks in the brain, longterm cultures are necessary. Most neuronal networks require between 21 - 28 days in culture to mature and will until that point undergo continued axonal outgrowth, fasciculation, and eventually synaptic pruning(8, 21). Furthermore, the way in which topographical features and modular organization affect the functional connectivity and signal propagation within engineered neuronal networks is still largely unexplored. As such, there is a need for experimental platforms which reliably and longitudinally support the study of dynamic structure-function relationships in engineered *in vitro* neuronal networks with controllable afferent-efferent connectivity during prolonged periods of times.

In this study, we use geometrical constraints inspired by the Tesla valve design to control afferent-efferent connectivity of rat cortical neurons. The Tesla valve was originally designed by Nikola Tesla more than 100 years ago to allow fluids to preferentially flow in a single direction without the assistance of any moving mechanical parts(22). The geometry of the design promotes liquid flow in the forward direction, while the liquid is looped back on itself when moving in the backward direction to effectively create an impenetrable barrier. This valvular conduit works in tandem with the laminar flow present in microfluidics, achieving highly efficient control of liquid flows(23). Here, we show that the same design can be used to establish neuronal networks with definable connectivity. We evaluate the structure and functions of networks during a culture period of 4 weeks. Our approach facilitates a robust method for the recapitulation and study of the development of complex topological structures of the brain in both health and disease. Additionally, implementation of the Tesla valve promotes highly efficient liquid separation between the neuronal populations, which can be useful in a range of applications, such as disease modelling and drug screening.

## Materials and Methods

### Design of Experimental Platform

Designs for the microdevices were created using Clewin 4 (WieWeb Software, Enschede), and are shown in **Figure S2.** Fabricated devices can be seen in **Figure S3.** The microfluidic platforms comprised two 5mm wide and 60 μm high compartments, henceforth referred to as nodes, connected by 20 microchannels, each 700 μm long, 10 μm wide and 5 μm high. The compartments employed a partially open design, to allow experimental control and reproducibility(24, 25). To promote unidirectional axonal growth, geometrical constraints inspired by the Tesla valve design were incorporated in the microfluidic channels. Furthermore, axon traps were incorporated on the postsynaptic side to misguide any outgrowing axons(16). A pattern of 4 μm diameter pillars with 4 μm interspacing was positioned on the presynaptic side to prevent neuronal somata from entering the channels. 100 μm pillars were also incorporated along the edges of the chambers to prevent the collapse of the 60 μm high compartments in front of the microtunnels. For the two-nodal microfluidic devices interfaced with microelectrode arrays (n = 5), 59 electrodes of 50 μm diameter were positioned evenly spread across the chips, of which 10 electrodes were placed selectively in the channels to confirm active connections between the neuronal populations. A reference electrode was split between the two chambers. Platforms with a single chamber interfaced with microelectrode arrays were used as controls (n = 6), in which the total seeding area was equal to the two-nodal devices. An experimental timeline is shown in **Figure 1.**

**Figure 1.**
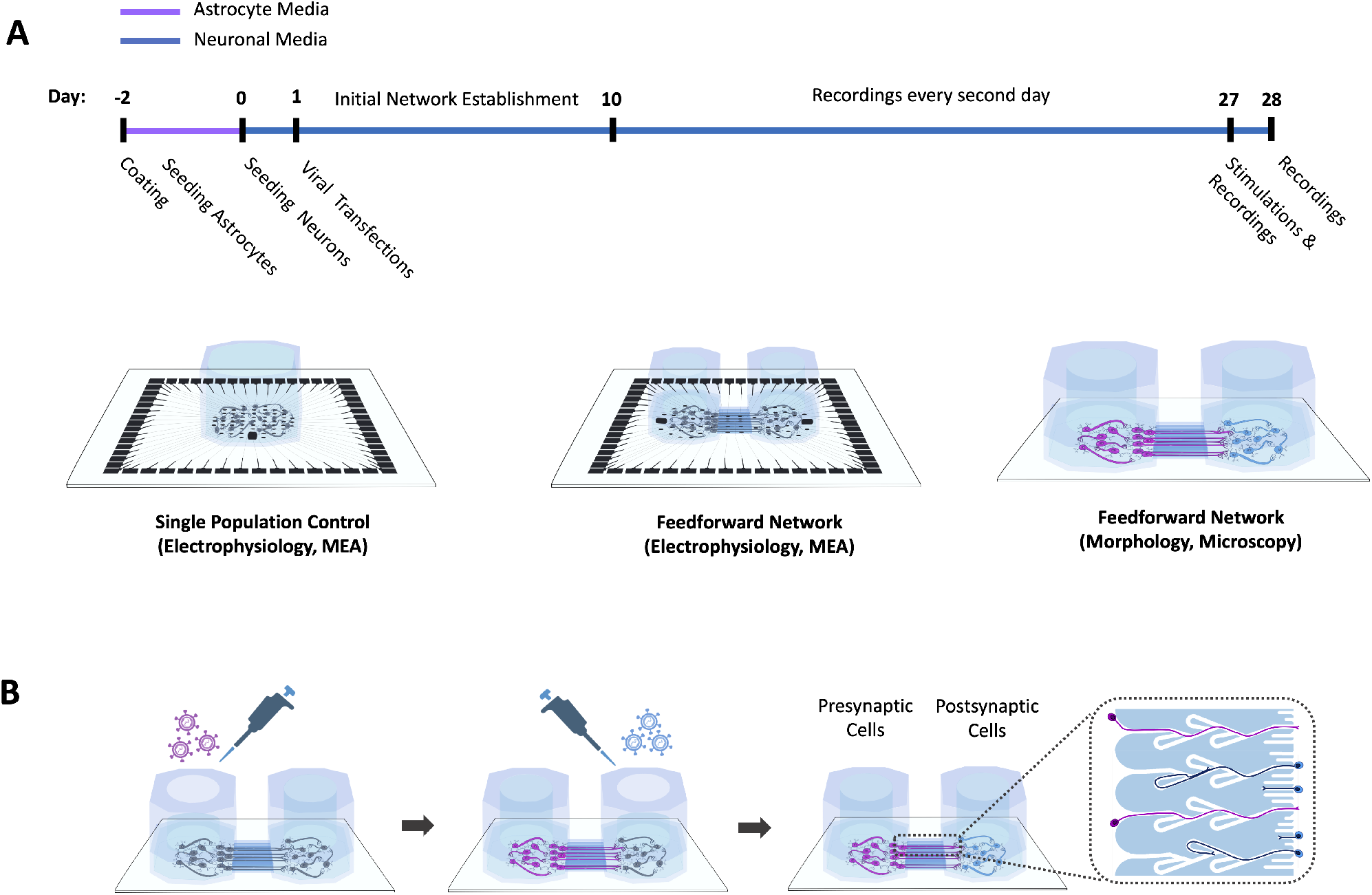
Experimental timeline. **(A.)** Neurons were grown in PDMS microdevices attached either to glass coverslips or custom-made microelectrode arrays. Elecrophysiological recordings were conducted every second day from day 10 to day 28 *in vitro* (DIV). Recordings after electrical stimulations were conducted at 27 DIV. **(B.)** On day 1, neurons grown in microfluidic chips attached to the glass coverslips were transfected with AAV 2/1 with a CMV promoter to induce ubiquitous expression of different fluorescent proteins in the two cell populations. This facilitated the examination of the presence of axons from the pre- and postsynaptic populations within the microchannels. The schematics are not to scale.

### Fabrication of Microfluidic MEAs

A schematic of the full fabrication process can be found in the supplementary materials (**Figure S1)**.

#### Mould for Microfluidic Chips

A soft lithography mould was made by application of a photoresist pattern on a 4-inch silicon wafer (Siltronix). The wafer was washed consecutively in acetone and isopropanol (IPA) for 1 min each to remove organic contaminants, followed by a plasma clean for 5 min in 100 sccm O_2_ plasma at 20 kHz generator frequency (Femto Plasma Cleaner, Diener Electronics). A 5 min dehydration bake was conducted at 120 °C to remove moisture. The photoresist mr-DWL5 (micro resist Technology GmbH) was spin-coated onto the substrate in two steps of 3000 rpm for 35 s at 1000 rpm/min and 4000 rpm for 2 s at 2000rpm*/*min, respectively, to achieve a final thickness of 5.0 μm (spin150, SPS-Europe B.V.). A soft bake was subsequently conducted, starting at 50 °C for 2 min before gradually increasing the temperature to 90 °C (5 °C min^-1^) where it was kept for 2 min. The wafer was left to cool down on the hotplate (5 °C min^-1^), before being left at room temperature (RT) to rest for 10 min. A maskless aligner (MLA150, Heidelberg) was used to transfer the MEA design onto the resist with a 405 nm laser at 300 mJ/cm^2^. After exposure, the film was post exposure baked equivalently to the soft bake, before being left to rest at RT for 1 h. The film was subsequently developed in mr-Dev 600 (micro resist Technology GmbH) for 120 ± 15 s before thoroughly rinsed in IPA. The substrate was thereafter hard baked at 120 °C for 30 min.

A second layer of photoresist mr-DWL40 (micro resist Technology GmbH) was spin-coated onto the substrate in two steps of 1200 rpm for 30 s at 500 rpm/min and 3500 rpm for 5 s at 1000 rpm/min, respectively, to achieve a final thickness of 60 μm. A soft bake was consecutively conducted, starting at 50 °C for 5 min before gradually increasing the temperature to 90 °C (5 °Cmin^-1^) where it was kept for 10 min. The wafer was left to cool down on the hotplate (5 °Cmin^-1^), before being left to rest at RT for 1 h. The film was exposed with a 405 nm laser at 600mJ/cm^2^. After exposure, the film was post exposure baked equivalently to the soft bake, before being left to rest at RT overnight. The film was subsequently developed in mr-Dev 600 (micro resist Technology GmbH) for 300 ± 15 s before thoroughly rinsed in IPA. The substrate was hard baked at 120 °C for 30 min. A profilometer (Dektak 150, Veeco) was used to confirm the intended heights. The finalized mould was eventually silanized (trichloro(1H,1H,2H,2H-perfluorooctyl(97 %)), Sigma-Aldrich) in vacuum for 2 h.

#### Microfluidic Chips

Silicon elastomer and curing agent (SYLGARD®184 elastomer kit, Dow Corning) was mixed, degassed, and cast in the photoresist mould at a ratio of 10:1. The PDMS was cured in an oven (TS8056, Termaks) at 65 °C for 4 h. The PDMS was peeled from the mould, and cell compartments were cut out with a 4mm diameter puncher. PDMS debris was removed using scotch tape, and the chips subsequently washed in acetone, 96 % ethanol and DI water for 1 min each. The chips were eventually left to dry overnight.

#### Microelectrode Arrays

The protocol for fabrication of microelectrode arrays was adapted and partly modified from van de Wijdeven *et al*.(26). 1mm thick 4-inch borosilicate wafers (100mm Borofloat33, Plan Optik) were used as substrates for the MEAs. The wafers were washed subsequently in acetone and IPA for 1 min each to remove organic contaminants, before being plasma cleaned for 5 min in 100 sccm O_2_ plasma at 20 kHz generator frequency. A 2 min dehydration bake was conducted at 100 °C to remove moisture. The photoresist ma-N 440 (micro resist Technology GmbH) was spin coated onto the substrates at 3000 rpm for 42 s at 500rpm/min for a final thickness of 4 μm (spin150, SPS-Europe B.V.). The film was left to rest for 10 min before being soft baked at 95 °C for 5 min. A maskless aligner (MLA150, Heidelberg) was used to transfer the MEA design onto the resist with a 405nm laser at 1800mJ/cm^2^. After exposure, the resist was developed in ma-D332/s (micro resist Technology GmbH) for 90 ± 10 s before thoroughly rinsed in DI water. Prior to e-beam evaporation, the substrates were descummed in 100 sccm O_2_ plasma for 1 min at 20 kHz generator frequency. An adhesive layer of 50 nm titanium was evaporated onto the substrates at 5 Å s^-1^, followed by 100 nm platinum at 2 Å s^-1^ (E-beam Evaporator, Pfeiffer Vacuum Classic 500). A lift-off process was subsequently conducted in acetone before the wafers were rinsed in IPA.

Prior to passivation layer deposition, another descum was conducted for 1 min. Next, a 470 nm thick insulation layer of silicon nitride (Si_3_N_4_) was deposited on the substrates using plasma enhanced chemical vapour deposition at 300 °C for 30 min consisting of 20.0 sccm SiH_4_, 20.0 sccm NH_3_ and 980 sccm N_2_ gas (PlasmaLab System 100-PECVD, Oxford Instruments). A second lithographic step was conducted using ma-N 440 according to the same protocol as previously described to define an etch mask. Inductively coupled plasma with 50.0 sccm CHF_3_, 10.0 sccm CF_4_ and 7.0 sccm O_2_ gas for 6.5 min was subsequently used to dry etch the silicon nitride above the electrodes and contact pads (Plasmalab System 100 ICP-RIE 180, Oxford Instruments).

A thin layer of platinum black was eventually electrodeposited on the electrodes (PP-Type Wafer Plating Electroplating Laboratory System, Yamamoto). The electrolyte bath consisted of aqueous 2.5 mmol chloroplatinic acid (H_2_PtCl_6_, 8wt% H_2_O, Sigma-Aldrich, 262587), in which the wafers were partially submerged using a custom-built wafer holder so that the liquid only covered the electrodes and not the contact pads (illustrated in **Figure S1B**). A Red Rod REF201 Ag/AgCl electrode (Hatch) was used as reference, and a platinized titanium plate as counter electrode. A paddle agitator was used at 60.0 rpm to reduce diffusion thickness, and the temperature was kept constant at 30 °C by an external temperature controller. A Palmsens 4 potentiostat (Palmsens) was used in chronoamperometric mode with a constant voltage of −0.4V for 3 min for all depositions. Eventually, the substrates were diced into 48×48 mm square MEAs with a wafer saw (DAD323, DISCO). The photoresist masks were subsequently stripped off in acetone, followed by a rinse in 96 % ethanol and DI water. To remove hardened photoresist and oxidize the top few nanometres of the silicon nitride layer into silicon dioxide (SiO_2_), the surface was plasma cleaned for 10 min in 160 sccm O_2_ at 32 kHz generator frequency.

#### Assembly of Microfluidic MEA Platforms

The microfluidic chips were bonded irreversibly to either MEAs or precleaned (acetone, 96 % ethanol and DI water) glass coverslips (24×24 mm Menzel-Gläser, VWR International). Both microfluidic chips and MEAs/coverslips were plasma cleaned with O_2_ plasma for 1 min in 200 sccm O_2_ plasma at 40 kHz generator frequency. Directly after, the PDMS chips were pushed gently onto the MEAs/coverslips. To facilitate alignment of the microfluidic chips to the MEAs, two drops of 70 % ethanol were placed in between the PDMS chip and the MEA, and alignment was performed under a stereomicroscope (Nikon SMZ800). For both coverslips and MEAs, bonding was finalized on a hotplate at 100 °C for 1 min followed by 5 min at RT under gentle pressure. To remove traces of ethanol from the platforms, three subsequent washes in DI water were conducted with 10 min intervals. Finally, the chips were filled with DI water to maintain hydrophilicity and sterilized in UV light overnight in a biosafety cabinet.

### Cell Culturing and Staining

#### Coating of Culturing Platforms

After sterilization, DI water was replaced by 0.1 mg/ml Poly-L-Ornithine solution (PLO) (Sigma-Aldrich, A-004-C) and the chips incubated at 37 °C and 5 % CO_2_ for 30 min. Subsequently, the PLO was replaced by fresh PLO, and the chips incubated at 37 °C and 5 % CO_2_ for another hour. Next, the chambers were washed three times with MQ water, before being filled up with laminin solution consisting of Leibovitz-15 Medium (Sigma-Aldrich, L5520) supplemented with 45.6 mg/ml sodium bicarbonate (Sigma-Aldrich, S6014) and 16 μg/ml natural mouse laminin (Gibco™, 23017015) and incubated at 37 °C, 5 % CO_2_ for 30 min. Subsequently, the laminin solution was replaced with fresh laminin solution, and the chips incubated for another 2 h. During all coating steps, a hydrostatic pressure gradient was created between the chambers by filling one chamber fuller than the other to assure proper flow of coating solution through the microfluidic channels.

#### Cell Seeding and Maintenance

Laminin solution was replaced by astrocyte media consisting of DMEM, low glucose (Gibco™, 11885084) supplemented with 15% Fetal Bovine Serum (Sigma-Aldrich, F9665) and 2% PenicillinStreptomycin (Sigma-Aldrich, P4333). Rat astrocytes (Gibco™, N7745100) were consecutively seeded at a density of 100 cells/mm^2^, i.e., 2000 cells per microchamber in the two compartmental devices and 4000 cells in the single compartmental devices. After two days of expansion, the astrocyte media was replaced with neuronal media consisting of Neurobasal Plus Medium (Gibco™, A3582801) supplemented with 2% B27 Plus (Gibco™, A358201), 1% Gluta-Max (Gibco™, 35050038) and 2% Penicillin-Streptomycin (Sigma-Aldrich, P4333). Additionally, Rock Inhibitor (Y-27632 dihydrochloride, Y0503, Sigma-Aldrich) was added at a concentration of 0.1% during seeding to increase survival rate. Pen-strep was not included in the microfluidic chips used for viral transfections until 4 DIV to avoid potential interference with transfection efficacy. Rat cortical neurons from Sprague Dawley rats (Gibco, A36511) were seeded at a density of 1000 cells/mm^2^, equalling 20 000 cells per chamber in the two-nodal platforms and 40 000 cells in the controls. Half the cell media was replaced with fresh cell media 4 hours after seeding, and again after 24 h. From here on, half the cell media was replaced every second day until the cultures were ended at 28 DIV.

#### Viral Transfections

To assess the efficacy of the microfluidic design at promoting unidirectional axonal growth, neurons in two-compartmental devices on glass coverslips were virally transfected with an AAV 2/1 serotype loaded with either pAAV-CMV-beta Globin intron-EGFP-WPRE-PolyA or pAAV-CMV-beta Globin intron-mCherry-WPRE-PolyA. This was done to ubiquitously express different fluorescent proteins in the two neuronal populations, aiding longitudinal monitoring of the structural connectivity in the microchannels during development. Transfections were started one day after seeding. A hydrostatic pressure gradient was used during the transfection to hinder viral leakage between the two nodes. 3/4 of the medium in the presynaptic chamber was removed, and AAV 2/1 viruses with a CMV promoter encoding GFP were added at a titer of approximately 5*e*^2^ viruses/cell (1*e*^7^ viruses per chamber). The cells were incubated at 37 °C and 5% CO_2_ for 3 h. Next, the presynaptic chamber was filled up with fresh neuronal medium, and 3/4 of the medium in the postsynaptic chamber was removed. AAV 2/1 viruses with a CMV promoter encoding mCherry were furthermore added to the postsynaptic chamber at a titer of 5*e*^2^ viruses/cell. The postsynaptic chamber was subsequently filled all the way up again after 3 h. The devices were finally left unperturbed for 48 h. While a small amount of cross-contamination of viruses was observed across the chambers, the described approach was found to yield the best trade-off between transfection efficacy and cross-contamination. Viral vectors were prepared in-house at the Viral Vector Core Facility, NTNU.

Images were acquired using an EVOS M5000 microscope (Invitrogen) using DAPI (AMEP4650), CY5 (AMEP4656), GFP (AMEP4651) and TxRed (AMEP4655) LED light cubes and Olympus UPLSAP0 4X/0.16 NA and 20x/0.75 NA lenses. Post-processing of images was conducted in ImageJ with the Fiji plugin to change the selected colour channels and add scale bars.

#### Immunocytochemistry

For immunocytochemistry, cells were seeded in μ-slide 8 Wells (Ibidi, GmbH, 80826) following the same protocol as for the microfluidics. Fixation was done using glyoxal solution based on the protocol by Richter *et al*.(27). The solution consisted of 71% MQ water, 20% ethanol absolute (Kemetyl, 100%), 8.0% Glyoxal solution (Sigma-Aldrich, 128465) and 1% acetic acid (Sigma-Aldrich, 1.00063). Cell medium was replaced with the fixative and the cells incubated at RT for 15 min, before washing the chambers with phosphate-buffered saline (PBS, Sigma-Aldrich, D8537) 3 times for 5 min each. Permeabilization was conducted using 0.5% Triton-X (Sigma-Aldrich, 1086431000) diluted in PBS for 5 min, before the chambers were washed 2 consecutive times in PBS for 5 min. Next, blocking solution consisting of 5% goat serum (Abcam, ab7481) diluted in PBS was added to the chambers, and the chips were left to incubate at RT on a shaking table at 30 rpm for 1 h. The blocking solution was thereafter replaced by primary antibodies mixed with 5% goat serum in PBS. A full overview of primary antibodies and their respective concentrations are listed in Table 1. The chips were subsequently placed on a shaker table at 30 rpm at 4 °C overnight. The following day, secondary antibody solution was prepared consisting of 0.2% secondaries and 5% goat serum diluted in PBS. Prior to mixing, the secondary antibodies were centrifuged at 6000 rpm for 15min to remove precipitates. Next, the primary antibody solution was removed from the chambers, and the chambers washed three times with PBS for 5 min each. Subsequently, secondary antibody solution was added to the chambers, and the cells incubated at RT on a shaker table at 30 rpm for 3 h in the dark. After this, the secondary antibody solution was replaced by solution consisting of the nuclear stain Hoechst (Abcam, ab228550) at a concentration of 1/2000 diluted in PBS and incubated at a shaking table at RT for 30 min. Finally, the chips were once again washed three times in PBS for 5 min each, then once in MQ water for 5 min, before eventually being filled halfway up with MQ water for imaging.

**Table 1.** Antibodies and concentrations used for Immunocytochemistry

| Marker | Catalogue Number | Concentration |
| --- | --- | --- |
| β3-Tubulin | Ab78078 | 1/1000 |
| Doublecortin | Ab153668 | 1/200 |
| GABA | Ab86186 | 1/1000 |
| GAP-43 | Sigma-Aldrich 9264 | 1/500 |
| GFAP | Ab7260 | 1/1000 |
| MAP2 | Ab5382 | 1/5000 |
| NeuN | Ab279295 | 1/500 |
| Neurofilament Heavy | Ab4680 | 1/5000 |
| PSD95 | Ab13552 | 1/250 |
| Synaptophysin | Ab32127 | 1/500 |
| Tbr1 | Ab183032 | 1/100 |

*All antibodies were purchased from Abcam unless otherwise stated.*

Imaging was conducted using a Zeiss 800 Airyscan Confocal Laser Scanning Microscope (CLSM) with Zeiss Zen Blue software. The microscope was equipped with a LSM 800 Laser Module URGB with diode lasers 405nm (5mW), 488nm (10mW), 561nm (10mW) and 640nm (5mW). A Zeiss Axiocam 503 color camera, 2.8 Mpix, 38 fps (color) was furthermore used for image acquisition.

#### Sample Dehydration and SEM Imaging

For the full protocol, see the supplementary materials.

### Electrophysiology and Data Analysis

#### Electrophysiological Recordings

The average impedance of the electrodes was measured to 34.5 ± 10.9 kΩ in 0.1% PBS Solution using a MEA-IT60 System (Multichannel Systems). Recordings were conducted using a MEA2100 workstation (Multichannel Systems) at a sampling rate of 25000 Hz. A temperature controller was used to maintain temperature at 37 °C (TC01, Multichannel Systems). A 3D-printed plastic construct covered by a gas-permeable membrane was used to keep the cultures sterile during recordings. Prior to recordings, the networks were allowed to equilibrate for 5 min at the recording stage. Each recording lasted for 15 min, and recordings were always conducted on the day following media changes.

#### Electrical stimulations

Stimulations were performed at 27 DIV using a train of 60 spikes at ± 800mV (positive phase first) of 200 μs duration with an inter spike interval of 5 s. Prior to stimulation, a baseline recording of 5 min was conducted to identify the most active electrode close to the centre of the chamber measured in number of spikes/s, which was subsequently chosen for stimulation. For the two-nodal platforms, the most active electrode in the postsynaptic node was stimulated first, followed by the most active electrode in the presynaptic node.

#### Data Analysis

All data analysis, including filtering and spike detection, was conducted in Matlab R2021b using a combination of borrowed and custom-made scripts. Raw data was filtered using a 4th order Butterworth bandpass filter, removing high frequency noise (above 3000 Hz) and low frequency fluctuations (below 300 Hz). A notch filter was used to remove 50 Hz noise caused by the mains hum. Both filters were run using zero-phase digital filtering with the Matlab function filtfilt to avoid changes in the relative position of the detected spikes. Spike detection was conducted using the Precise Timing Spike Detection (PTSD) algorithm developed by Maccione *et al*.(28) The threshold was set to 11 times the standard deviation of the noise, the maximum peak duration to 1 ms and the refractory time to 1.6 ms.

Burst detection was subsequently performed using the logISI approach developed by Pasquale *et al*.(29), with a minimum of 4 consecutive spikes set as a threshold for a burst to be detected. The hard threshold for the interspike interval both within and outside of the burst core was set to 100 ms. This number was chosen based on the original manuscript reporting the method(29), as well as manual inspection of the burst detection performance. For analysis of spontaneous network activity, network bursts were furthermore detected using the logIBEI approach(30), with at least 20% of all active electrodes in the network required to participate for the activity to be classified as a network burst. Network synchrony was measured as the median number of active electrodes participating in each network burst during a recording.

For each network burst, the summed-up spiking activity of each node was binned into 20 ms bins, and the resulting tuning curves of the two nodes were plotted. If a network burst was detected, but only one of the nodes had more than 30 detected spikes within the time span of the network burst, the network burst was evaluated as contained within that individual node. If more than 30 spikes were detected within both nodes, the network burst was considered to have propagated between the nodes. If the network burst was found to spread between the two nodes, the temporal position of the peaks of the two tuning curves were compared to evaluate whether the network burst spread from the pre-to the post-synaptic node or vice versa. Only response delays between 20 ms and 240 ms were included in the analysis as delays shorter or longer than this made it difficult to infer directionality or made it unlikely for the activity to be part of the same network burst, respectively. This delay time is also consistent with the propagation times reported by others for *in vitro* neuronal networks(31). A raster plot and plot of the binned firing rate showing the detection and analysis of the spread of networks bursts are shown in **Figures 2A** & **B**. Resulting tuning curves of a network burst spreading from the pre-to the postsynaptic node, and vice versa, are shown in **Figure 2C**.

**Figure 2.**
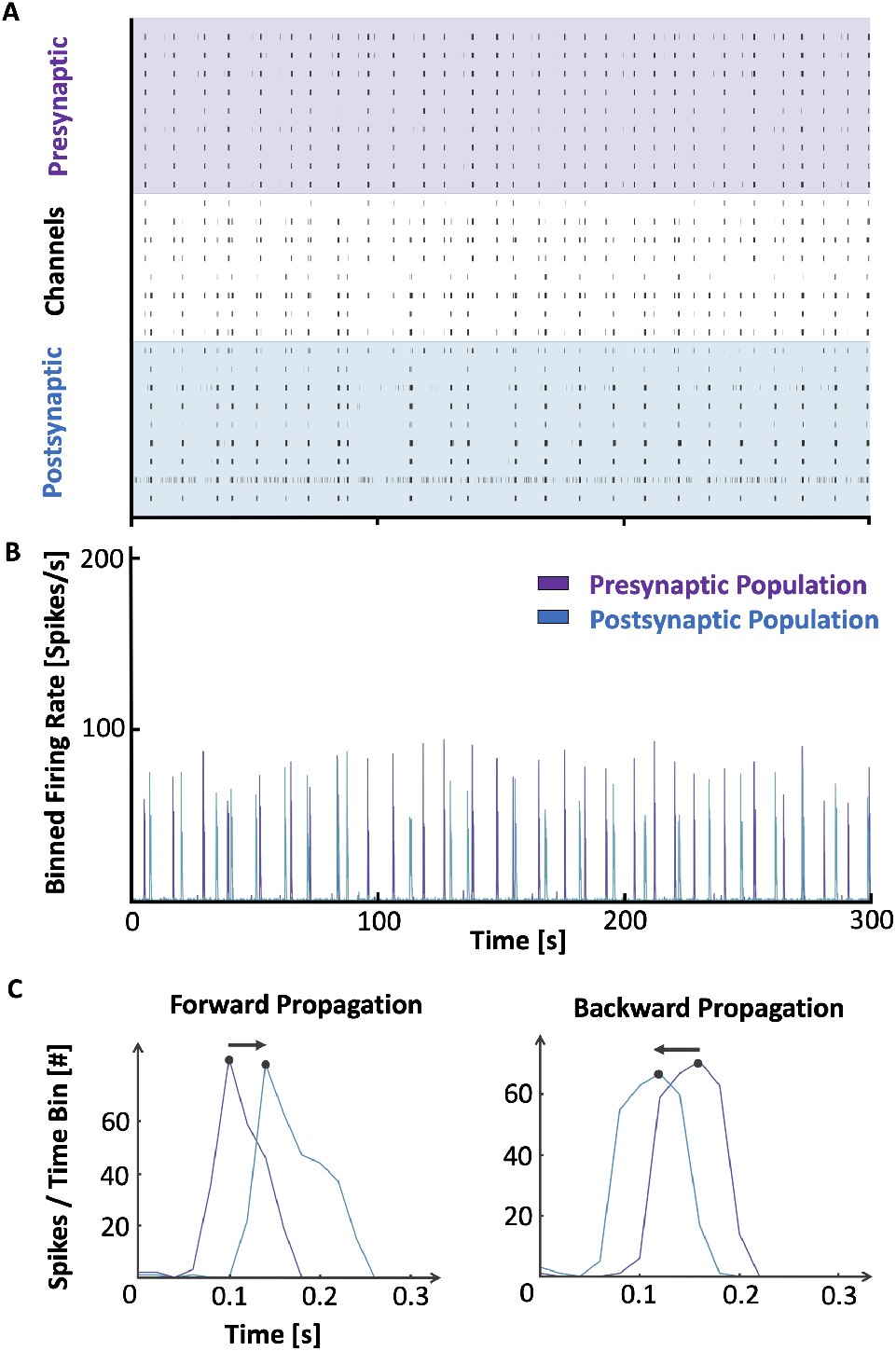
Illustration of analysis of network bursts in two-nodal feedforward networks. **(A.)** 300 s raster plot showing the synchronized propagation of multiple network bursts across the two nodes. **(B.)** Binned network activity showing the total firing rate in the two nodes during the same period of time as in A. **(C.)** Tuning curves of two individual network bursts, showing how the summed-up activity in both nodes were plotted and used to determine the direction of activity propagation.

Stimulation data was run through the SALPA filter developed by Wagenaar *et al*.(32), and each stimulated time point was followed by 15 ms blanking to avoid stimulation artifacts being detected as spikes. Peristimulus time histograms were made for each platform by binning the data in the 300 ms following a stimulation into 20 ms time bins, and the average response of the network in each bin for the 60 individual stimulations was plotted. Raster plots were made using a customized version of the SpikeRasterPlot function developed by Kraus(33). Heatmaps showing 20 ms binned network activity following stimulations were created using a customized version of the Matlab function heatmap developed by Deoras(34).

Spike data for the spontaneously evoked activity was binned into 50 ms time bins, and functional connectivity was evaluated using pairwise mutual information through the information theory toolbox developed by Timme & Lapish(35). Graph theoretical metrics were calculated using the Brain Connectivity Toolbox developed by Rubinov & Sporns(36). These were furthermore used to plot custom designed graphs showing node colour as spike frequency (spikes/s), node size as PageRank centrality, edge colour as the functional connectivity (pairwise mutual information) and node edge colour as the community calculated using the Louvain algorithm(37). Nodes were positioned according to the relative position of the electrodes on the actual microfluidic MEAs. Average graph metrics were also plotted over time.

For all plots, customized colormaps were created using the Matlab versions linspecer(38) and colorBrewer2(39), based on the web tool colorBrewer developed by Brewer *et al*.(40).

## Results

### Tesla Valve Inspired Design Promotes Unidirectional Connectivity Between Neuronal Nodes In Vitro

Expression of the fluorescent proteins from the viral transfections, used to assess the structural connectivity, was observed after 7 – 14 days in *vitro* (DIV). This allowed us to evaluate the efficacy of the Tesla valve microfluidic design at promoting unidirectional axonal outgrowth longitudinally as the networks matured. We found that the efficacy of the microdevice design in facilitating asymmetric axon growth could be tuned by the dimensions of the Tesla loops. A loop diameter of 40 μm, height of 60 μm and channel turning angle of 140° promoted directionality with primarily axons from the presynaptic node filling up the microchannels. Yet, as the networks matured and the channels filled up with axons from the presynaptic node, some axons from the postsynaptic node managed to grow in a zig-zag pattern within the Tesla loops to reach the opposite node (**Figure 3A**). As this was undesired, the loops were redesigned. Decreasing the loop diameter to 30 μm and increasing the height to 70 μm resulted in improved control of the afferent connectivity by directing the axons from the postsynaptic node back to the chamber of origin within the first few Tesla loops (**Figure 3B**). Controls with bidirectional microchannels were also tested to demonstrate the bilateral growth of axons in the channels in the absence of any geometrical constraints (**Figure S4)**.

**Figure 3.**
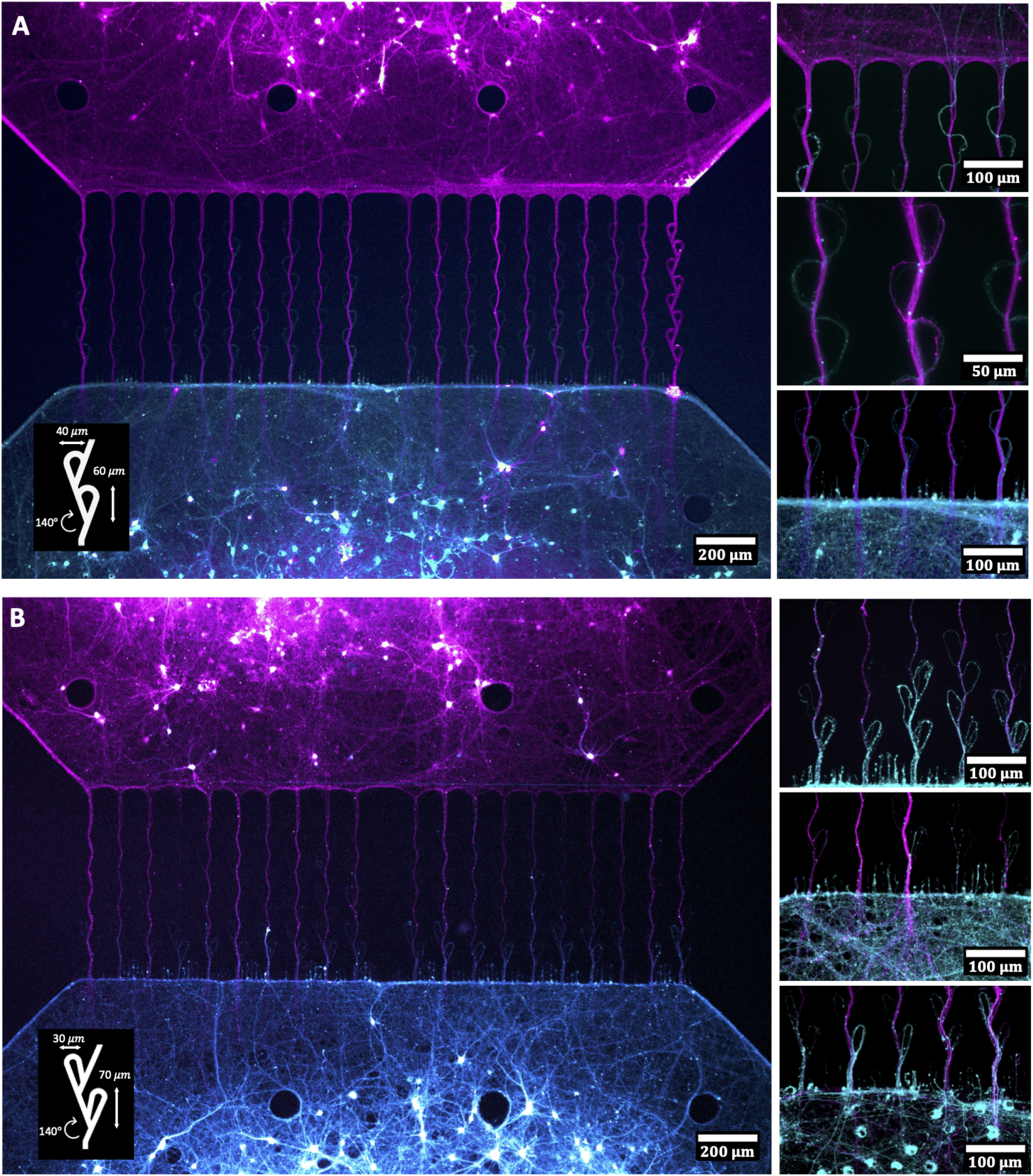
Images of the selective growth of axons through the Tesla valve inspired microfluidic chips. **(A.)** A loop diameter of 40 μm and height of 60 μm promoted directionality but still allowed for some postsynaptic axons to pass through the channels by exploiting the Tesla valves themselves. **(B.)** Decreasing the valve diameter to 30 μm and increasing the height to 70 μm effectively redirected the postsynaptic axons back to their chamber of origin within the first 3-4 valves. This effect was found to be stable throughout the culturing period. Black spots in the chambers indicate areas where 100 μm pillars were implemented in the design to keep the roof from collapsing in front of the microtunnels. Not all pillars were properly attached to the glass surface, hence the uneven number of black spots in the figures.

To further assess the robustness of our approach, geometries based on previously published designs were fabricated and compared with the Tesla valve design when using the same cell culture conditions (**Figure S6)**. Designs implementing trapping structures within the channels, similar to the one demonstrated by Gladkov *et al*.(12), were commonly found to cause extensive fragmentation of axons within the channels. This effectively caused an accumulation of cell debris within the trapping structures, furthermore hindering axons from the presynaptic population from reaching the opposite node, as well as confounding imaging. Designs redirecting the axons, such as the loop-back(16), were found to be more effective but still allowed multiple axons from the postsynaptic node to reach the presynaptic node, as illustrated in **Figure S6A**.

### Structural Maturation must be Considered when Evaluating Afferent-Efferent Network Connectivity

Immunocytochemistry performed at 7, 14 and 21 DIV confirmed a gradual change in the expression levels of developmental and cytoskeletal markers important for neuronal migration and growth. A selection of the images are shown in **Figure 4.** Doublecortin and Growth-Associated Protein 43 (GAP43) have both been found critical for neurodevelopment and growth cone migration, respectively, and were used as markers for immature neurons(41, 42). Both markers were found to be expressed until 21 DIV, indicating that the networks were still undergoing substantial axonal outgrowth, and thereby structural maturation at this point. The cytoskeletal markers Microtubule-Associated Protein 2 (MAP2) and *β*3-Tubulin start being expressed shortly after axonogenesis and continue to be expressed as the cells mature(43–47). These markers were used to assess structural network maturation. Both markers showed a high expression level throughout the network development, and indicated aggregation of neuronal somata and dense fasciculation already after 14 DIV (**Figure 4A**).

**Figure 4.**
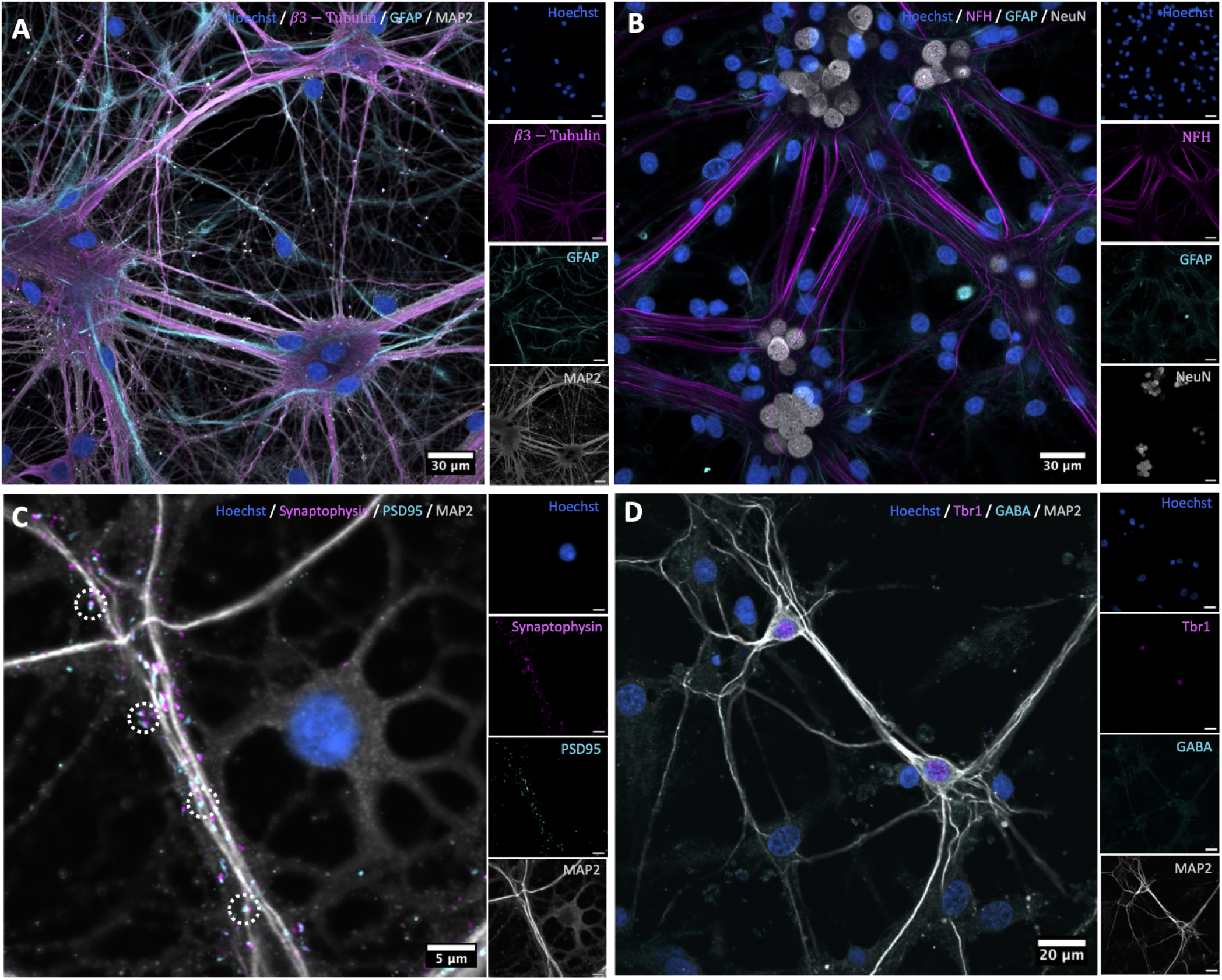
Immunocytochemistry. **(A.)** The neuronal networks exhibited high degree of clustering and axonal bundling after 14 DIV, demonstrated by the cytoskeletal marker proteins *β*3-Tubulin and MAP2. **(B.)** Expression of the mature markers Neurofilament Heavy (NFH) and NeuN indicating high degree of maturation after 21 DIV. **(C.)** Colocalization of the pre- and postsynaptic markers synaptophysin and PSD95 at 21 DIV indicating mature synapses. **(D.)** Expression of the glutamatergic receptor Tbr1 and GABA at 21 DIV indicating the presence of both excitatory and inhibitory activity.

The expression of Neuronal Nuclear Protein (NeuN) and Neurofilament Heavy (NFH) were used as markers for mature neurons(48, 49). Both markers were expressed at 21 DIV (**Figure 4B**). Images showing the development in expression profiles of the neuronal and astrocytic markers over time can be seen in the supplementary materials (**Figure S4)**. As can be clearly delineated from these images, fluorescence intensity of NFH was significantly higher at 21 DIV compared to 14 DIV. At 21 DIV, colocalization of synaptophysin and PSD95 also verified the presence of mature synaptic connections (**Figure 4C**)(50–52). The presence of glutamate receptors and GABA furthermore confirmed the presence of both excitatory and inhibitory activity in the network, known to be important for excitatory-inhibitory homeostasis and proper network functioning (**Figure 4D**)(53, 54). Staining for GFAP revealed a continued proliferation of astrocytes throughout network development for all cultures(55). These images confirm that the networks had started reaching structural maturity at 21 DIV, but were still undergoing structural changes even at this point as indicated by the high expression of GAP43.

### Spontaneous Activity Spreads in a Feedforward Manner Following Connectivity Establishment Between the Two Nodes

To evaluate the effect of the controlled afferent connectivity on network activity, neuronal cultures were grown on microfluidic chips interfaced with customized microelectrode arrays (MEA) to facilitate measurements of their electrophysiological activity (**Figures 5A** & **5B**). Given the high temporal resolution of spikes detected using MEA recordings, this approach was suitable both for assessing the functional connectivity within each node and for tracking signal propagation between them with high accuracy. To increase the signal-to-noise ratio of the noisy extracellular recordings in the open compartments, nanoporous platinum black was deposited on the MEA electrodes (**Figures 5C** & **5D**). Some electrodes were placed in the microchannels to follow signal propagation between the nodes (**Figures 5E** & **5F**).

**Figure 5.**
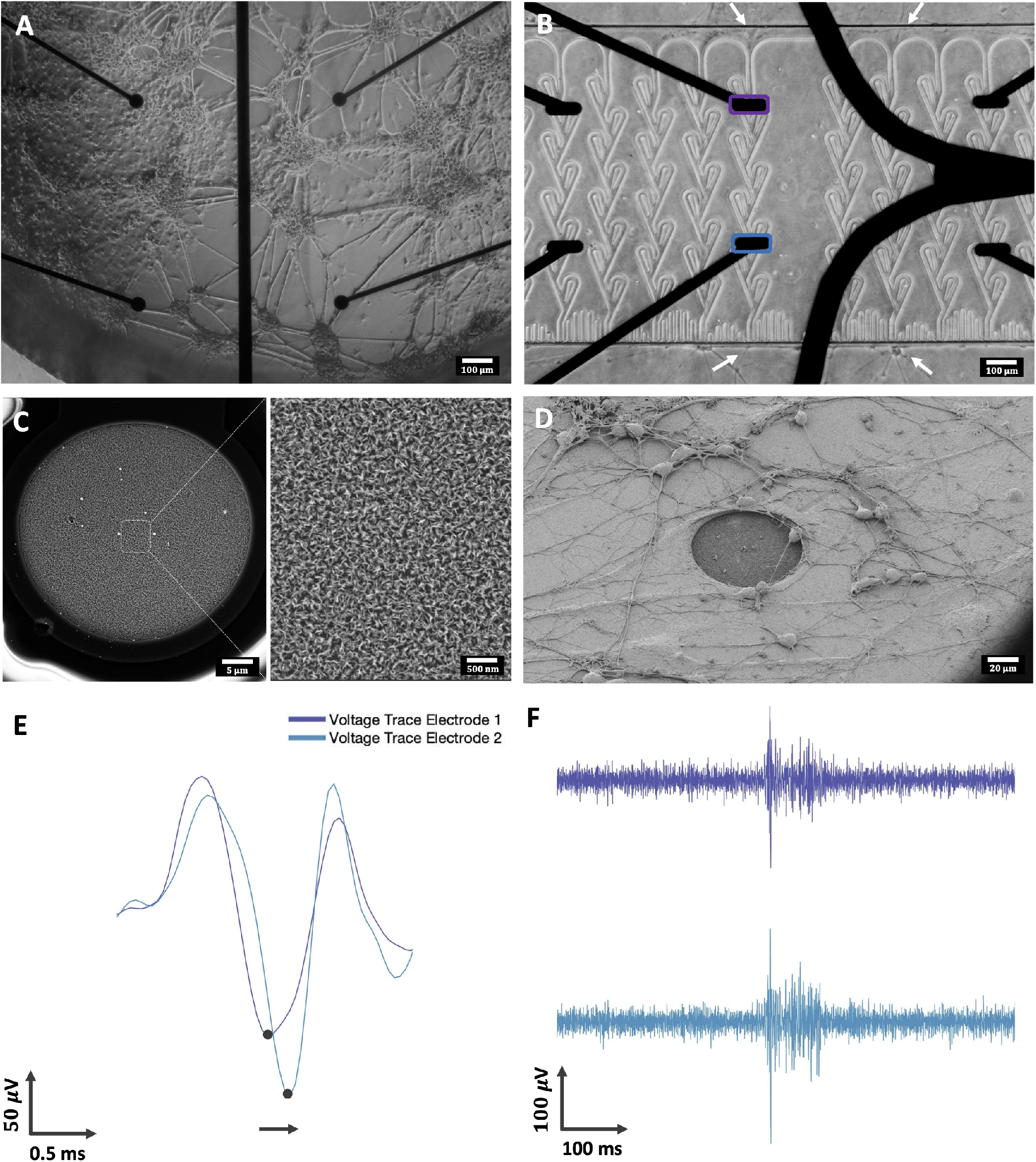
The microfluidic chips were interfaced with microelectrode arrays for electrophysiological recordings. **(A.)** Micrograph showing clustered networks growing on top four of the microelectrodes. **(B.)** Micrograph demonstrating neurite outgrowth through the microfluidic channels. **(C.)** SEM image of an individual electrode with deposited nanoporous platinum black. **(D.)** SEM image of neurons growing on top of a single microelectrode, demonstrating biocompatibility of the nanoporous electrodes. **(E.)** Voltage traces from two electrodes positioned along a microfluidic channel, demonstrating forward propagation of an action potential. **(F.)** Voltage traces of two electrodes positioned along a microfluidic channel showing the propagation of a single burst.

As individual spikes are challenging to track between the nodes, due to the high level of bursting activity, network bursts were consequently used to evaluate whether signals primarily moved from the pre-to the postsynaptic node. As detailed in the methods section, tuning curves of both nodes were created and their peaks were identified to evaluate the temporal spread of synchronized activity between them (see **Figure 2)**. When two nodes are unidirectionally connected, it is expected that signals will propagate from the pre-to the postsynaptic node, but not the other way around. The percentage of network bursts spreading from either the pre-to postsynaptic node, or vice versa, for each of the 2-nodal networks are shown with bars in **Figure 6A**. Graphs showing the corresponding total number of network bursts are shown in the same figures (secondary y-axis). Given the structural variability between the individual cultures, the activity in each microdevice was analysed separately. At DIV 10-12, nodes started exhibiting network-wide bursts. Bursts were initially found to establish independently and at a similar rate in both nodes in a single microdevice. At around three weeks *in vitro*, three out of five two-nodal unidirectionally connected networks (networks 1, 2 & 3) showed efficient and unidirectional transmission of network bursts from the pre-to the postsynaptic node. A few network bursts were analysed to have spread between the nodes also earlier during the network development. However, due to the high frequency of bursts, these could also have been localized network bursts in the two nodes overlapping in time without spreading between the nodes. For the cultures with the most prominent functional connectivity between the nodes, the proportion of feedforward network bursts thereafter increased for a few days until almost all global network bursts propagated from the pre-to the postsynaptic node. Box charts showing the delay times between the maximum response in the pre- and postsynaptic populations during network bursts for each of the five 2-nodal networks are included in **Figure S10.**

**Figure 6.**
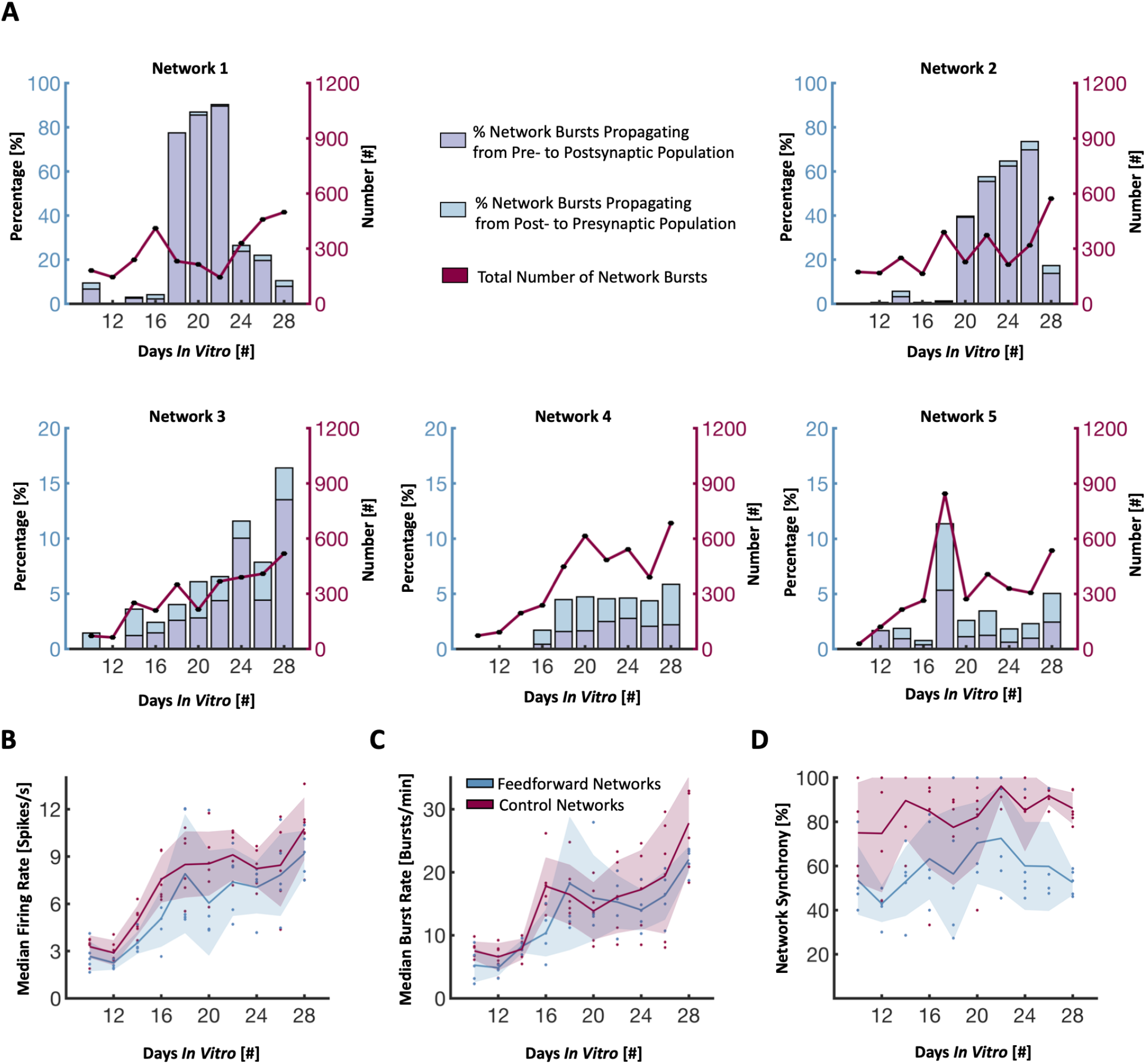
Development of spontaneous network activity. **(A.)** Histograms for each individual network showing the percentage of total network bursts being propagated either from the pre-to postsynaptic network, or vice versa, during the network development. The primary y-axis is scaled to 20% for networks 3-5. The total number of network bursts for each 15 min recording is shown along the secondary y-axis. Both firing rate **(B.)** and burst rate **(C.)** increased comparably over time for both the two-nodal networks and the controls. **(D.)** Network synchrony, measured as the percentage of the network participating in bursts, increased over time for both network types until around 3 weeks *in vitro*. After this, the synchrony decreased again towards 50% for the feedforward networks, indicating more localized activity in the two individual nodes.

The spike and burst rate of both the two-nodal networks and the single population controls were found to increase with time throughout the duration of the experiments (**Figures 6B** & **6C**), consistent with previous studies using *in vitro* neuronal cultures(29, 31, 56). Furthermore, the median percentage of the active electrodes taking part in network bursts, used as a measure of synchrony in the networks, are shown in **Figure 6D**. As can be seen from the plot, the control networks quickly moved towards 100%, indicating a high degree of synchrony across the networks. The two-nodal networks were initially close to 50%, consistent with network bursts being localized in the individual nodes, before increasing as the nodes in some networks connected and synchronized at around 21 DIV. Towards the end of the experiment, the synchrony in the two-nodal networks decreased again, as will be elaborated on in the discussion section.

### Electrical Stimulations Induce Feedforward Activity Propagation

Electrical stimulations were applied at 27 DIV to study signal propagation in the two-nodal microdevices. For two nodes that are unidirectionally connected, we expected evoked activity in the presynaptic node to spread to the postsynaptic node, but induced activity in the postsynaptic node to remain within that node. Stimulations were applied to the electrode showing the most spontaneous activity prior to stimulations in first the postsynaptic node, followed by the presynaptic node. The stimulations consisted of 60 consecutive 200 μs pulses of 800mV amplitude with a 5 s interspike interval. As expected, stimulations of the two-nodal networks with no apparent functional connectivity (networks 4 & 5) did not induce activity in the opposite neuronal node regardless of the node being stimulated. For the networks with spontaneous network bursts appearing to propagate between the nodes (networks 1, 2 & 3), stimulations of the presynaptic node induced a clear response in the postsynaptic node, confirming that the networks were functionally connected. Conversely, stimulations of the postsynaptic node did not induce any marked responses in the presynaptic node, as illustrated with a network burst in one of the networks in the raster plot in **Figure 7A**. The peristimulus time histograms showing the average response of one network are shown in **Figure 7B**, demonstrating a consistent forward propagation of network bursts in response to stimulations of the presynaptic node. **Figure 7C** shows a temporal heat map of a single stimulation in both the presynaptic (upper row) and postsynaptic (lower row) node for one network.

**Figure 7.**
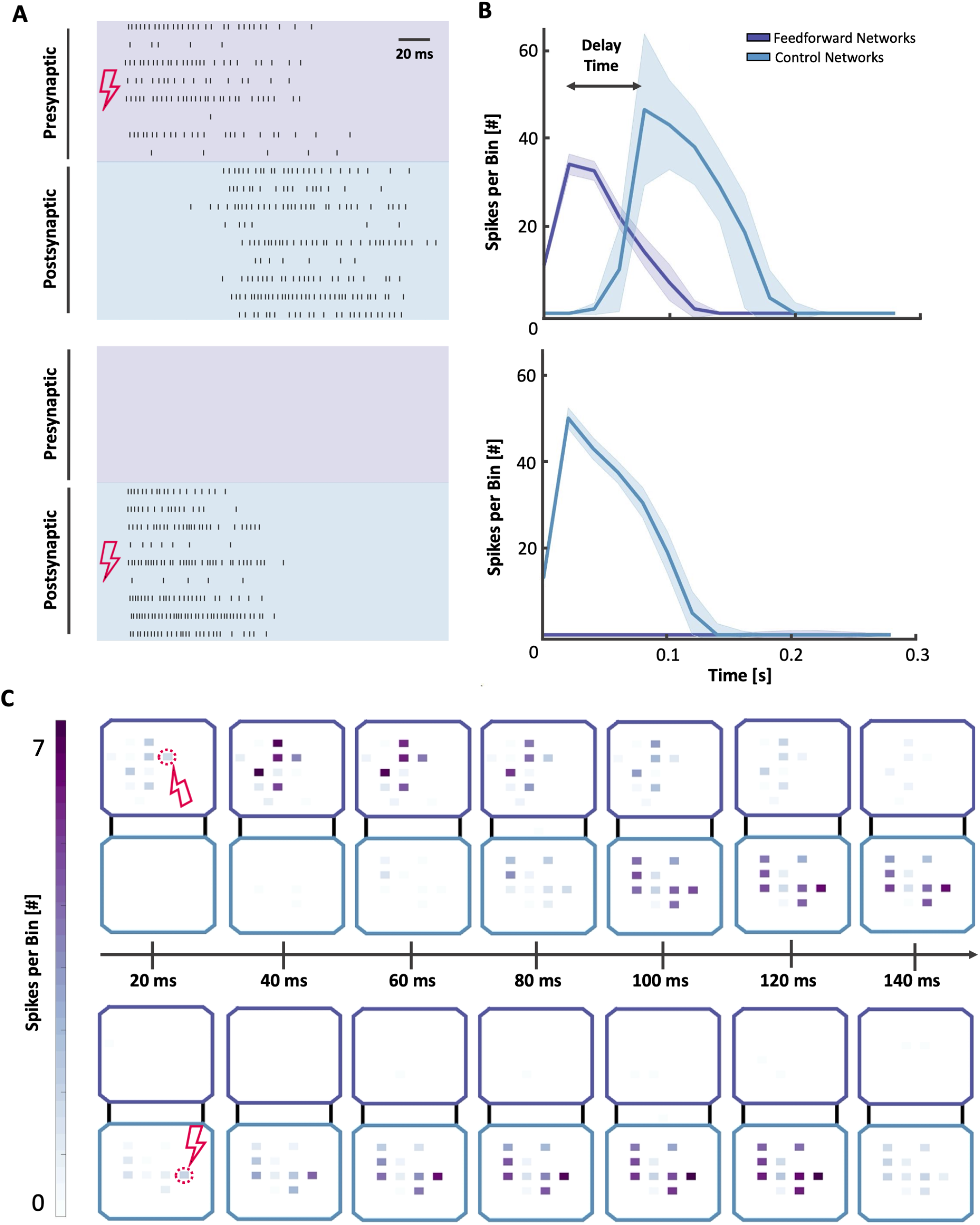
Illustration of response to electrical stimulations. **(A)** Raster plots showing propagation of a single network burst initiated by stimulation of the presynaptic and postsynaptic nodes, respectively. **(B)** Peristimulus time histograms showing the average response of 60 stimulations on each of the two nodes. Stimulations of the presynaptic node induced a clear response in the postsynaptic node, while no response was observed in the presynaptic node upon stimulation of the postsynaptic node. **(C)** Heat maps showing the binned (20ms) instantaneous firing of each active electrode in the 140ms following a stimulation in the pre- and postsynaptic nodes, respectively, further illustrating the spread of activity from the pre-to the postsynaptic node, but not vice versa.

### Graph Theoretical Analysis Indicates More Efficient Network Organization in the Two-Nodal Feedforward Network

Functional connectivity matrices and corresponding graphs were calculated based on pairwise mutual information, as detailed in the methods section. **Figures 8A** & **8B** show representative graphs of one two-nodal network and one control network, respectively. As can be seen from the graph representations, the nodes (here referring to individual measurement points) within the two-nodal populations could be effectively delineated into two separate communities, while the single population controls showed more random network organization with less distinguishable communities. While a network with only two communities is shown in **Figure 8B**, several of the control networks also had a higher number of communities with no apparent spatially ordered structure. This is furthermore supported by the plot of the maximized modularity in **Figure 8C**, showing a significant difference between the two-nodal networks and the controls. The difference in the global routing efficiency (**Figure 8D**) and characteristic path length (**Figure 8E**) of the two-nodal networks and the controls however appeared insignificant.

**Figure 8.**
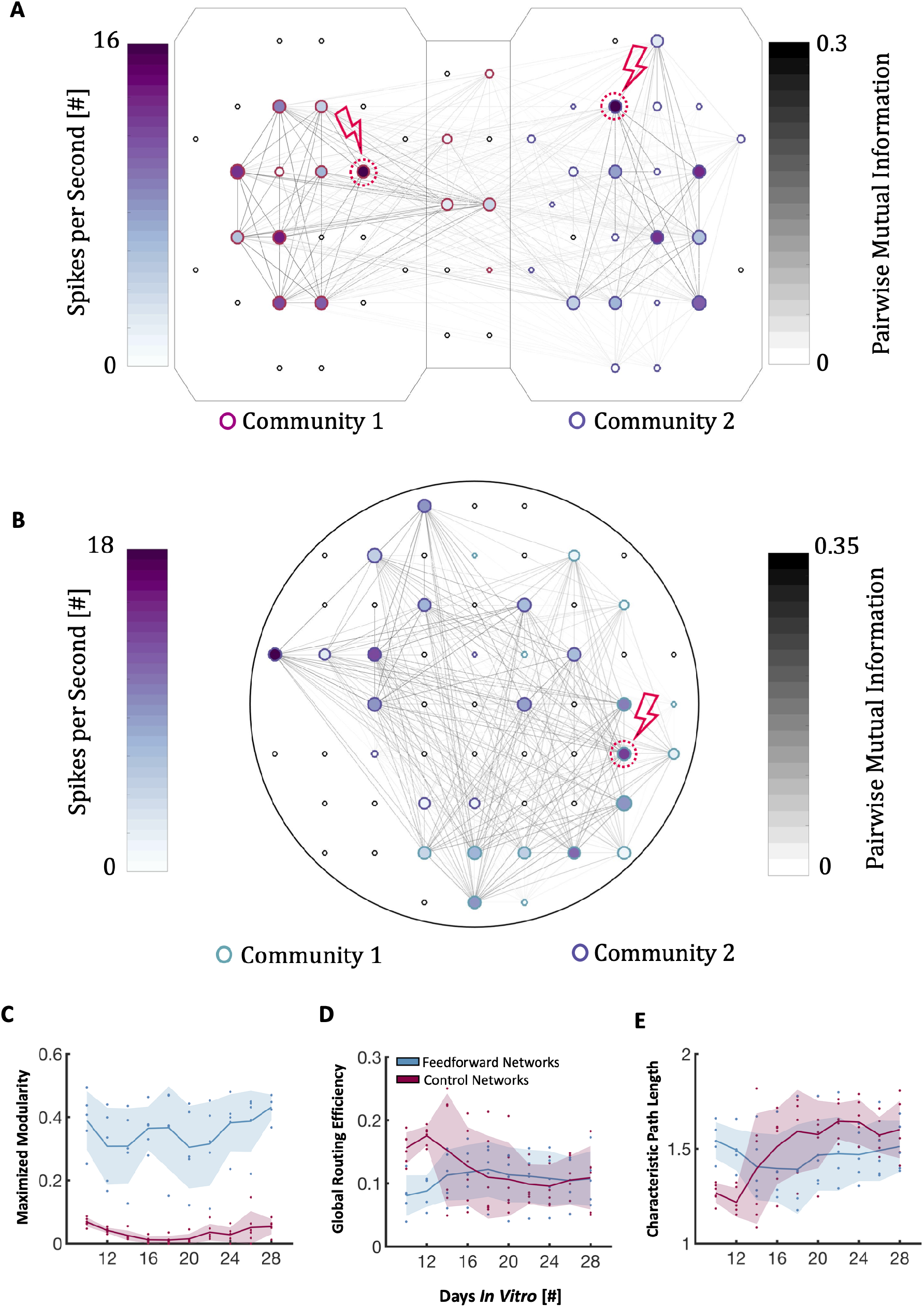
Graph Theoretical Analysis. **(A.)** Representative graph showing one of the two-nodal networks. Node colour represents firing frequency, edge colour the pairwise mutual information, node size the PageRank centrality and node edge colour the community belonging. **(B.)** Representative graph showing one of the control networks, demonstrating less clearly distinguishable modules. **(C.)** Plots showing that the maximized modularity is significantly higher for the two-nodal networks, while global routing efficiency and characteristic path length are more comparable in size.

## Discussion

In this study, we have explored the use of microfluidic devices with designs inspired by the Tesla valve to control the outgrowth of neurites between two neuronal populations. Our results indicate a high efficacy of these geometries at establishing controllable afferent connectivity both structurally (**Figure 3)** and functionally (**Figures 6** & **7**). Another key advantage of our microdevice is the ability to longitudinally assess both the structure and function of these networks, enabling sound analysis of dynamic structure-function relationships. Most studies looking at network development within microfluidic platforms with geometrically constrained microtunnels perform structural and/or functional analysis as early as 14 DIV, with 21 DIV being the maximum observation window reported in the literature. In our study, we observed these networks for a minimum of 28 DIV, and provide evidence that the networks still undergo structural (**Figure 4)** and functional (**Figure 6)** development up to this point. Long-term monitoring, i.e., at least 28 DIV, so that the networks can reach maturity, is therefore essential to evaluate how the microfluidic designs and embedded topographies influence network establishment and activity.

While only three out of five microdevices showed uniquivocal functional connectivity, it is worth noting that the channels used in this study were rather long (700 μm). Previous studies have shown that the dendritic arborization of rat cortical neurons can grow to above 300μm in culture(57), and that a microfluidic design of at least 400 to 450 μm is necessary to hinder dendritic branches from crossing the channels(10, 58). Using the GFP/mCherry to label neurites, we were able to show that the Tesla valve design effectively redirected nearly all neurites within three to four valves (equalling a distance of around 300 μm). With eight valves used in this study, a reduction to four valves might thus still be sufficient to control the afferent connectivity but could promote more efficient growth through the channels and thereby increase the percentage of cultures reaching functional connectivity. This could potentially also reduce the required time for the networks to reach functional connectivity between the nodes.

As the number of feedforward network bursts increased in some of the cultures, as shown in **Figure 6,** the generation of synchronous activity within the postsynaptic node (i.e., the number of network bursts contained within the postsynaptic node without spreading to the presynaptic node) seemed to decrease substantially. During the initial establishment of structural connections between the two nodes, it is likely that the newly attained afferent input of the postsynaptic node from the presynaptic node caused network depolarizations which effectively synchronized the activity of the two nodes over time. After a few days, the occurrence of feedforward burst propagation seemed to decrease again and the cultures appeared to go back to a state in which most network bursts were contained within the separate nodes. While the decrease in transmission of activity between the nodes might alternatively suggest decreased functional connectivity as a result of axonal degeneration in the microchannels or synaptic pruning, electrical stimulations of the presynaptic node still initiated activity that readily propagated between the nodes (**Figure 7)**. Furthermore, images of transfected neuronal populations, as the ones shown in **Figure 3,** demonstrate that the structural integrity of most of the cultures used for structural analysis was retained until the end of the experimental period at 28 DIV.

Notably, the modularity increased again after 20 DIV (**Figure 8C**), indicating more localized activity in the chambers. This is consistent with the bursts ceasing to propagate between the chambers in **Figure 6B**. Such a trade-off between functional separation and integration has previously been shown to enhance the functional capacity and input representation of modular networks(59). This is also in line with computational studies showing that modularity is a key determinant for the complex dynamics of cortical information processing, and that the coexistence of both segregated and integrated activity is beneficial to catalyse activity that does not immediately die out, but concurrently does not give rise to synchronized events spreading across the entire network(60–62). In support of this, previous studies of *in vitro* neuronal networks have indicated that physically constraining neuronal networks into modular configurations might be beneficial to increase the gating capacity and temporal coordination between different parts of a neuronal network(63, 64). Additionally, it could potentially increase the dynamical repertoire of activities in the networks by reducing the global synchronization(65).

It is also worth mentioning that the relative scarcity in the propagation of network bursts in two of the cultures (network 4 & 5) does not necessarily rule out any forms of structural or functional connectivity between the two nodes. In a previous study by Pan *et al*., it was estimated that a minimum of 100 axons were necessary to reliably transmit a burst from one population to another(7). With 20 microchannels connecting the two populations in our microdevices, on average 5 axons would have to cross each channel to reach this level of structural connectivity. As shown in **Figure 3,** as much as 15 – 20 individual protrusions can be seen entering the post-synaptic chamber from some of the channels. Based on the findings in the study by Pan *et al*., this level of connectivity should therefore in principle render the networks capable of reliably transmitting bursts between the nodes. However, whether these protrusions were individual axons or collaterals from the same axons is difficult to deduce. Additionally, it is possible that neurites do not grow as readily on the Si_3_N_4_ used as an insulation layer on the microfluidic MEAs as compared to glass due to surface functionalization characteristics, which could possibly explain why not all the two-nodal chips achieved synchronized activity. However, as proposed by Yamamoto *et al*., the highest dynamical richness might be reached at the verge of physical disconnection. Whether transmission of network bursts is necessary for a node to exert influence over another node is thus not given, and it has also previously been suggested that synchronized activity preferentially flows within a single neuronal population even in a well-connected two-nodal culture(66).

In fact, the function of network bursts is still not fully understood. It is widely accepted that they present an important feature of cultured neuronal networks(67, 68). They have also been shown to have a rich repertoire of activity patterns(31, 68), and to be able to represent and extend information from external stimuli(69). However, some reports have claimed that network bursts signify a form of developmental arrest in *in vitro* networks due to their lack of external input(67). Others have proposed that these bursts exhibit patterns resembling the slow wave oscillations characterizing sleep *in vivo*(70), or that they might play a role in erasing parts of the networks short-term memory(69). It has also been experimented with reducing such global network events either electrically(67), pharmacologically(70, 71), or chemogenetically(72), to understand their function and increase the computational complexity of the network. In our study, the global spread of network bursts rendered them useful for evaluating the unidirectional spread of information, and hence to demonstrate the efficacy of our model at establishing structured, unidirectionally connected neuronal assemblies.

In our study, we used a two-nodal design, but this can easily be scaled up to include more nodes, enhancing the topological complexity and hence the relevance of the platform when modelling neuronal circuits. As the connection strength between the subpopulations is also highly dependent on the number of axons crossing the microtunnels, experimenting with the channel length and number could provide valuable insights into the modulation of connection strengths on network activity dynamics in future studies(7). Such increased complexity could potentially bring further insights into how different nodes exert influence over one another, and hence give rise to richer repertoires of activities better resembling the diverse patterns of activity, input representations and adaptations characterizing the complex computational dynamics seen *in vivo*.

## Conclusion

*N*euronal assemblies in the brain are connected through highly specific topological projection sequences and hierarchies aiding efficient, directional information processing and computations. Being able to recapitulate microscale and mesoscale aspects of this organization with neuroengineering approaches would enable the establishment of more robust model systems for studying structure-function dynamics of *in vitro* neuronal networks. In this study, we have presented a novel microdevice for controlling the afferent connectivity between two neuronal nodes *in vitro* using microchannels with geometries inspired by the Tesla valve design. Furthermore, by combining these devices with viral tools to fluorescently label the neuronal populations in combination with embedded nanoporous microelectrode arrays for functional analysis, we demonstrate the power of our model system for longitudinal studies of structure-function dynamics of such modular neuronal networks. Our results indicate that the networks still undergo significant structural alterations after 3 weeks *in vitro*, and that the number of network bursts spanning both nodes increases over time. Furthermore, structurally connected nodes readily propagate activity evoked by electrical stimulations in a feedforward fashion. Potential modifications such as reducing the channel lengths might further improve growth of axons through the channels without compromising unidirectionality. This technology can be utilized to build model systems resembling the mesoscale architectural motifs found in the brain to further advance our understanding of how such networks organize, compute, adapt and represent information in both health and disease.

## AUTHOR CONTRIBUTIONS

The author contributions follow the CRediT system. **NWH**: Conceptualization, Methodology, Software, Investigation (chip design & manufacturing, cell experiments, electrophysiology, formal analysis), Writing – Original Draft, Visualization. **ÅBT**: Investigation (immunocytochemistry and confocal microscopy), Writing – Review & Editing. **PS, AS, IS**: Conceptualization, Methodology, Writing – Review & Editing, Funding acquisition.

## ACKNOWLEDGEMENTS

NTNU Enabling technologies and Health Mid-Norway are acknowledged for funding this research. The Research Council of Norway is acknowledged for the support to the Norwegian Micro- and Nano-Fabrication Facility, NorFab, project number 295864.

We would like to thank Dr. Rajeevkumar Nair Raveendran at the Viral Vector Core Facility, Kavli Institute for systems neuroscience, for designing and preparing the AAV2/1 viruses used for structural analysis, Prof. Michela Chiappalone and Prof. Sergio Martinoia, University of Genova for generously providing the scripts for the Precise Timing Spike Detection algorithm and the logISI burst detection, and Edevard Hvide MSc for acquisition of the SEM image in **Figure 2D**. We acknowledge Senior Engineer Astrid Bjørkøy and the Center for Advanced Microscopy (CAM) at the Department of Physics, Faculty of Natural Sciences, NTNU for technical assistance and access to confocal microscopy infrastructure.

## Supplementary Materials

**Figure S1.**
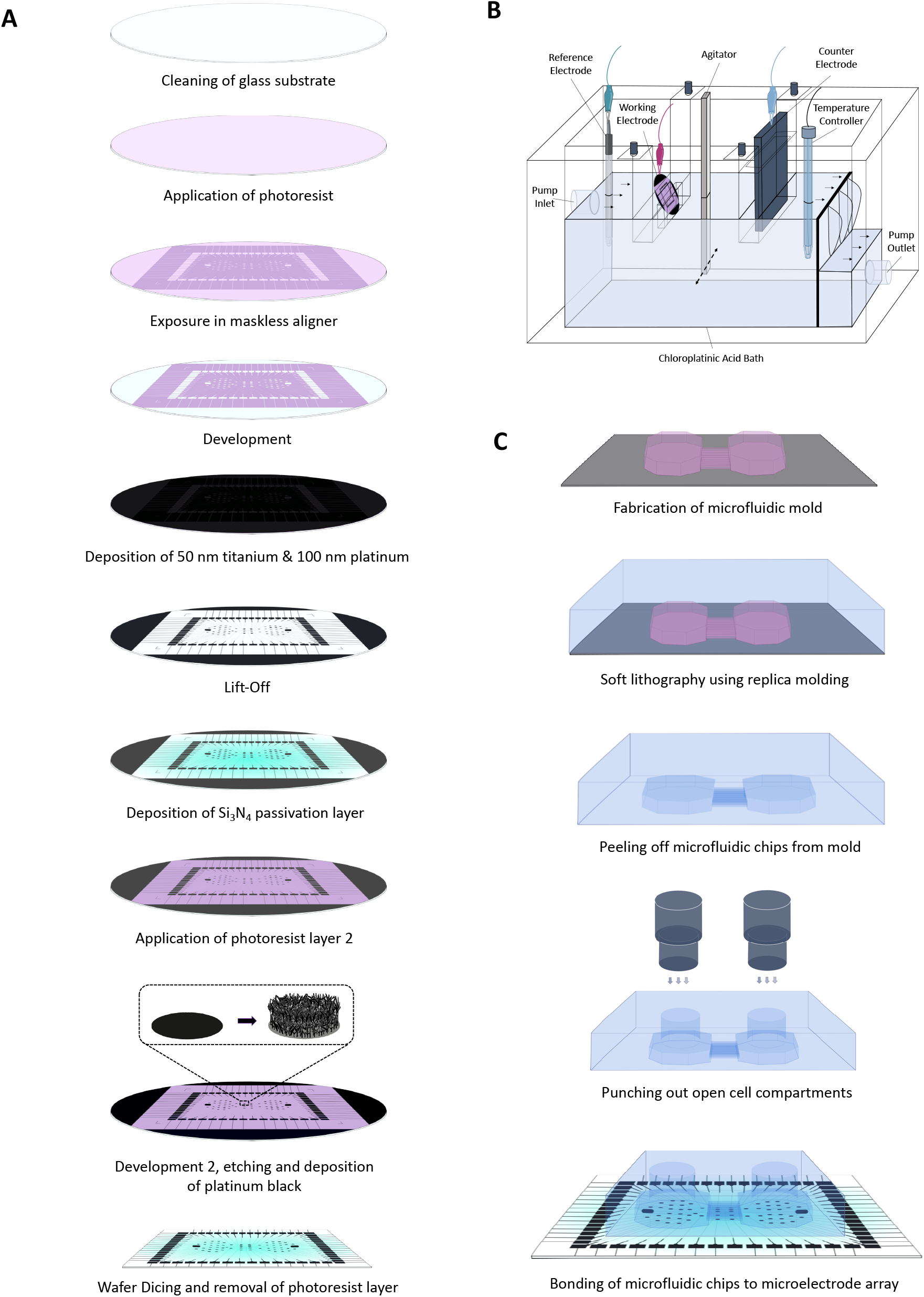
Schematics of fabrication process, as described in the methods section of the main paper. **(A.)** Fabrication of microelectrode arrays. **(B.)** Setup for electrochemical deposition of nanoporous platinum black. **(C.)** Fabrication of microfluidics chips.

**Figure S2.**
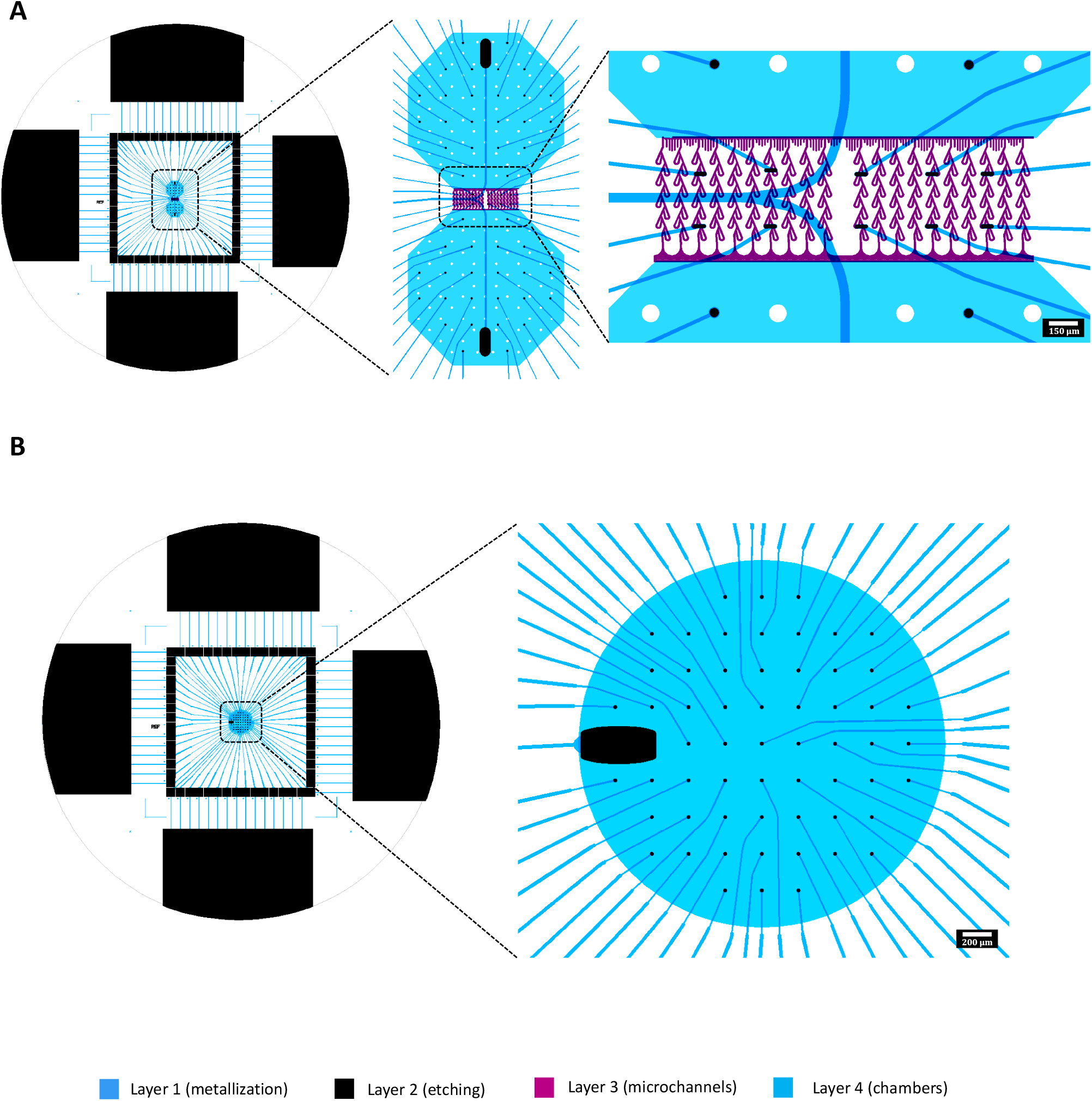
CAD designs of **(A.)** the two-nodal microfluidic platform with Tesla valve microfluidic design and **(B.)** the single population control platform. Layer 1 is the design for metallization, layer 2 for the etch mask, layer 3 for the microfluidic channels and layer 4 for the cell compartments.

**Figure S3.**
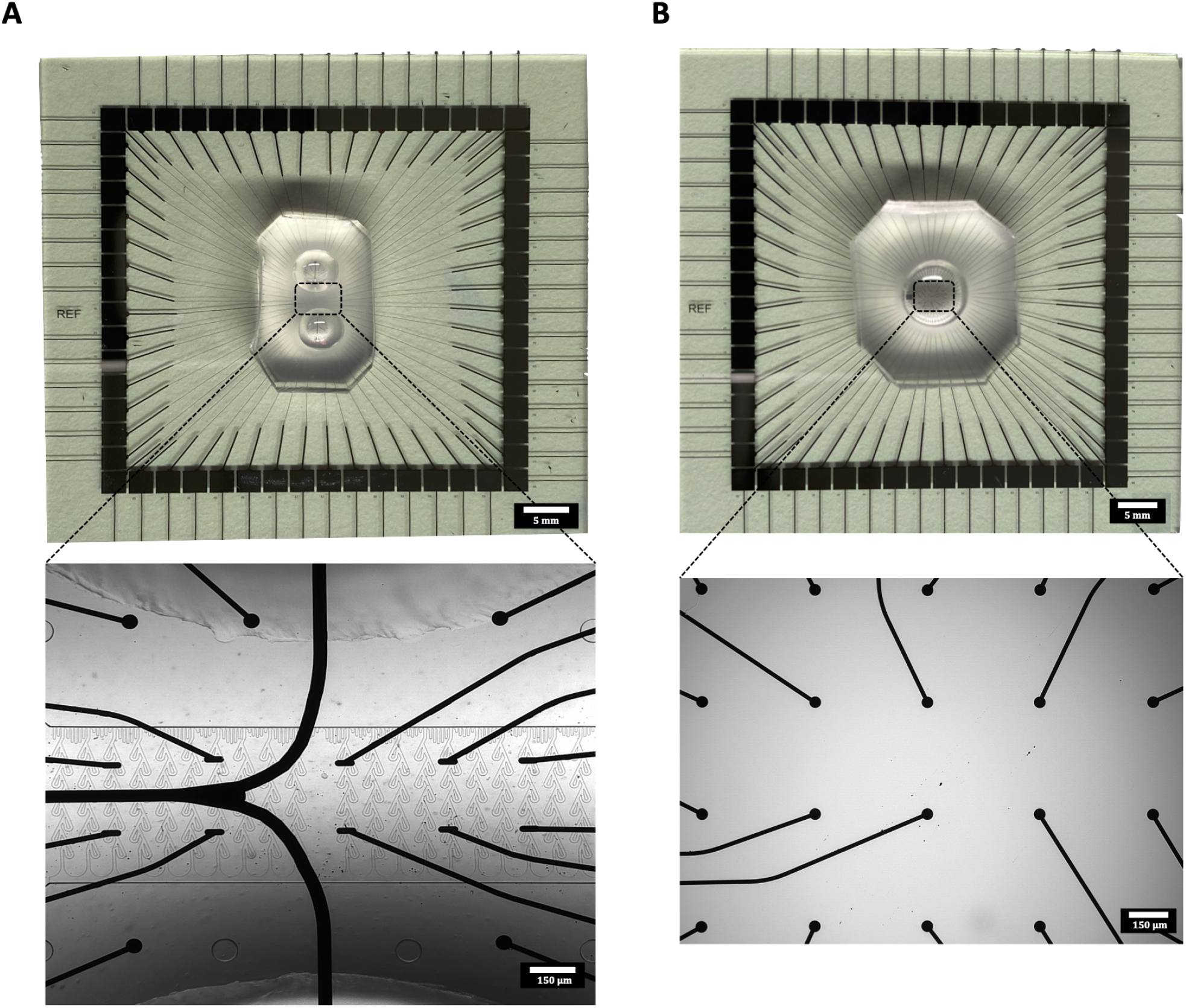
Images showing the fabricated microfluidic chips bonded to nanoporous microelectrode arrays. **(A.)** A two-nodal microfluidic device with Tesla valve microfluidic channels. **(B.)** A control platform with a single well for culturing cells.

**Figure S4.**
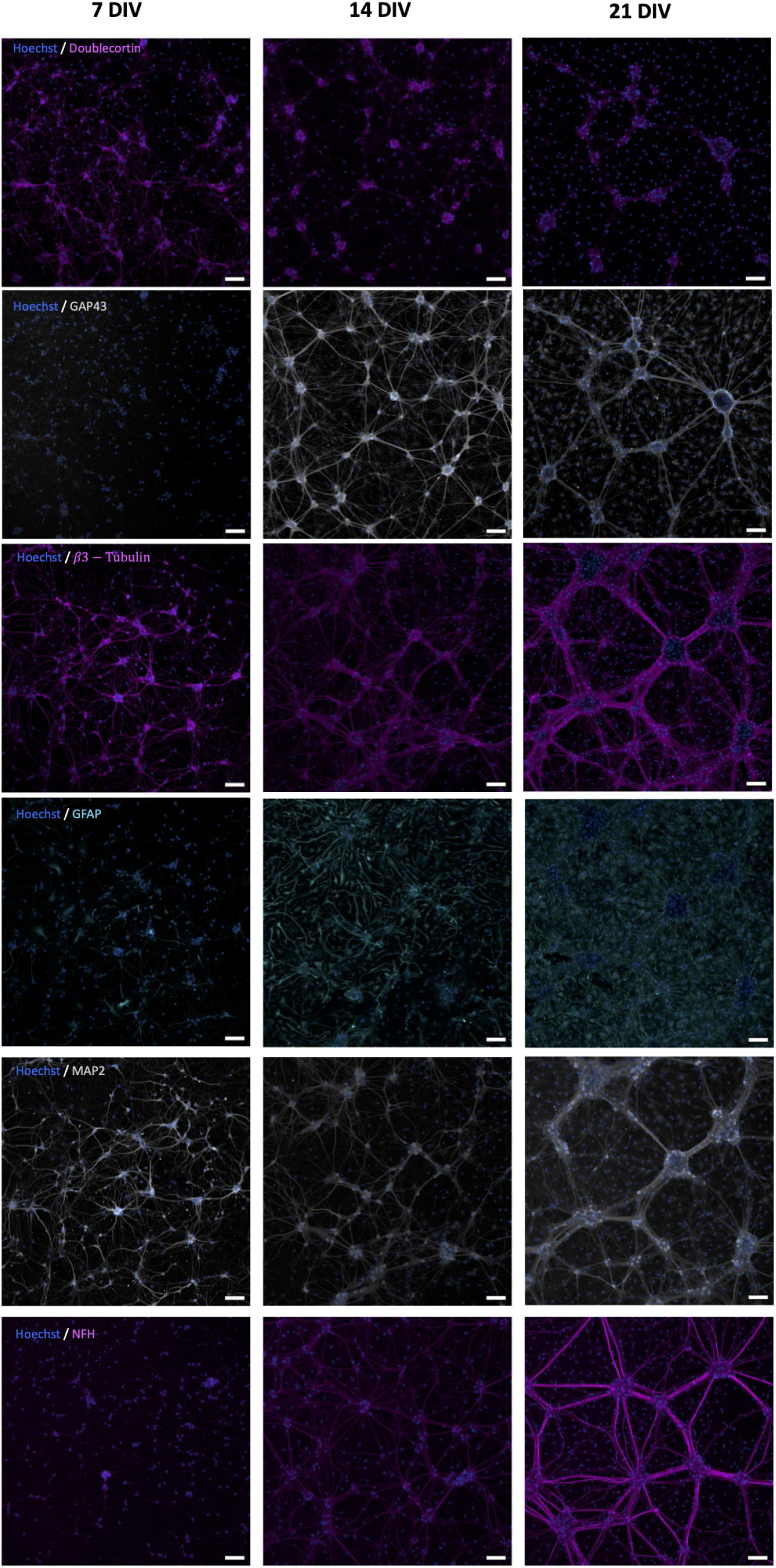
Immunocytochemistry showing longitudinal expression of doublecortin and GAP43 (neurodevelopmental markers), *β*3-Tubulin and MAP2 (cytoskeletal markers), NFH (mature cytoskeletal marker) and GFAP (astrocytic marker). Scale bars: 30μm.

**Figure S5.**
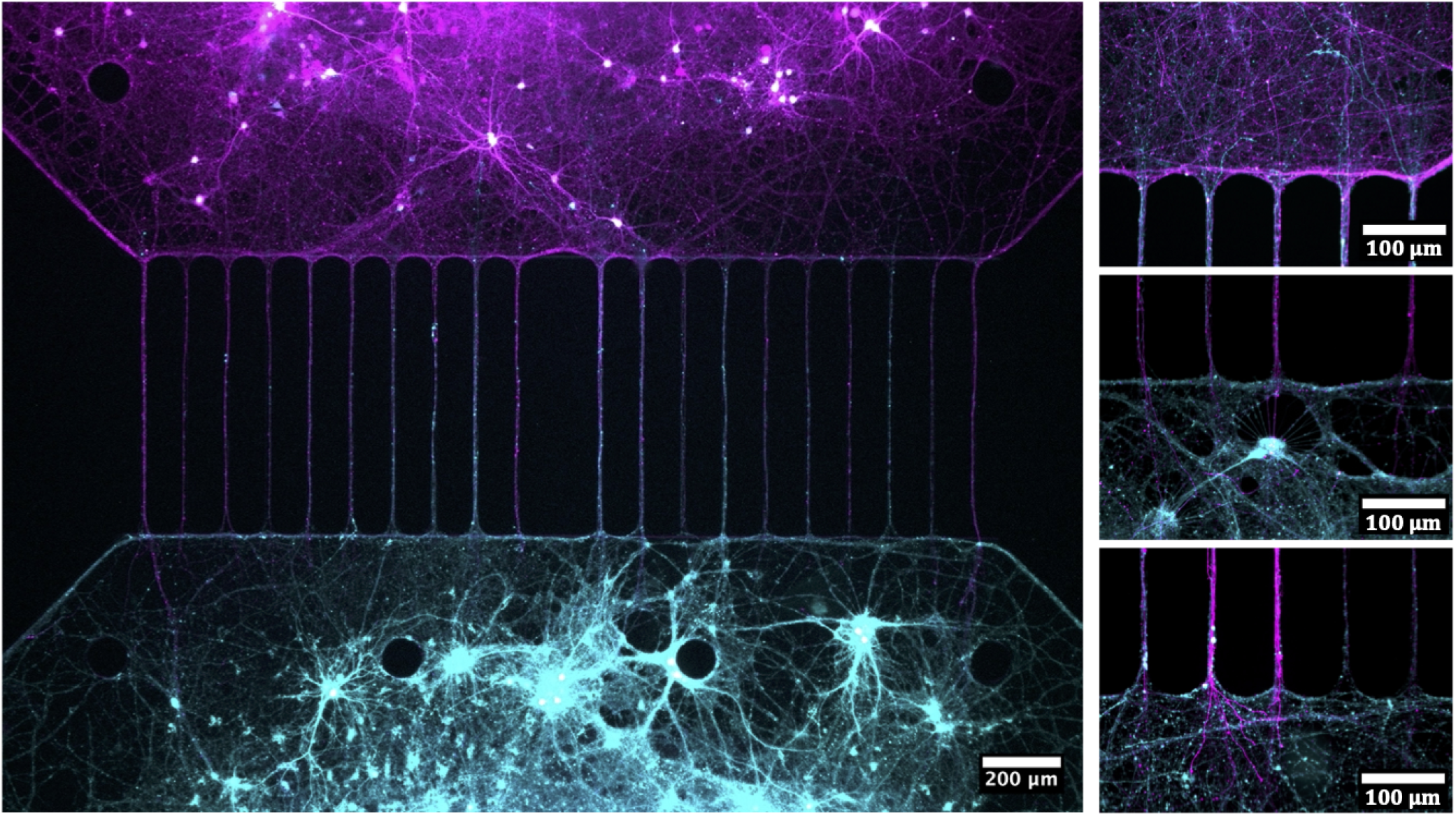
Neuronal cultures transfected with AAV2/1 with a CMV ubiquitous promoter for expression of GFP/mCherry in microfluidic controls with bidirectional channels. The figure illustrates the clear bidirectional growth in the channels when no geometrical constraints are used to promote unidirectional axonal growth.

**Figure S6.**
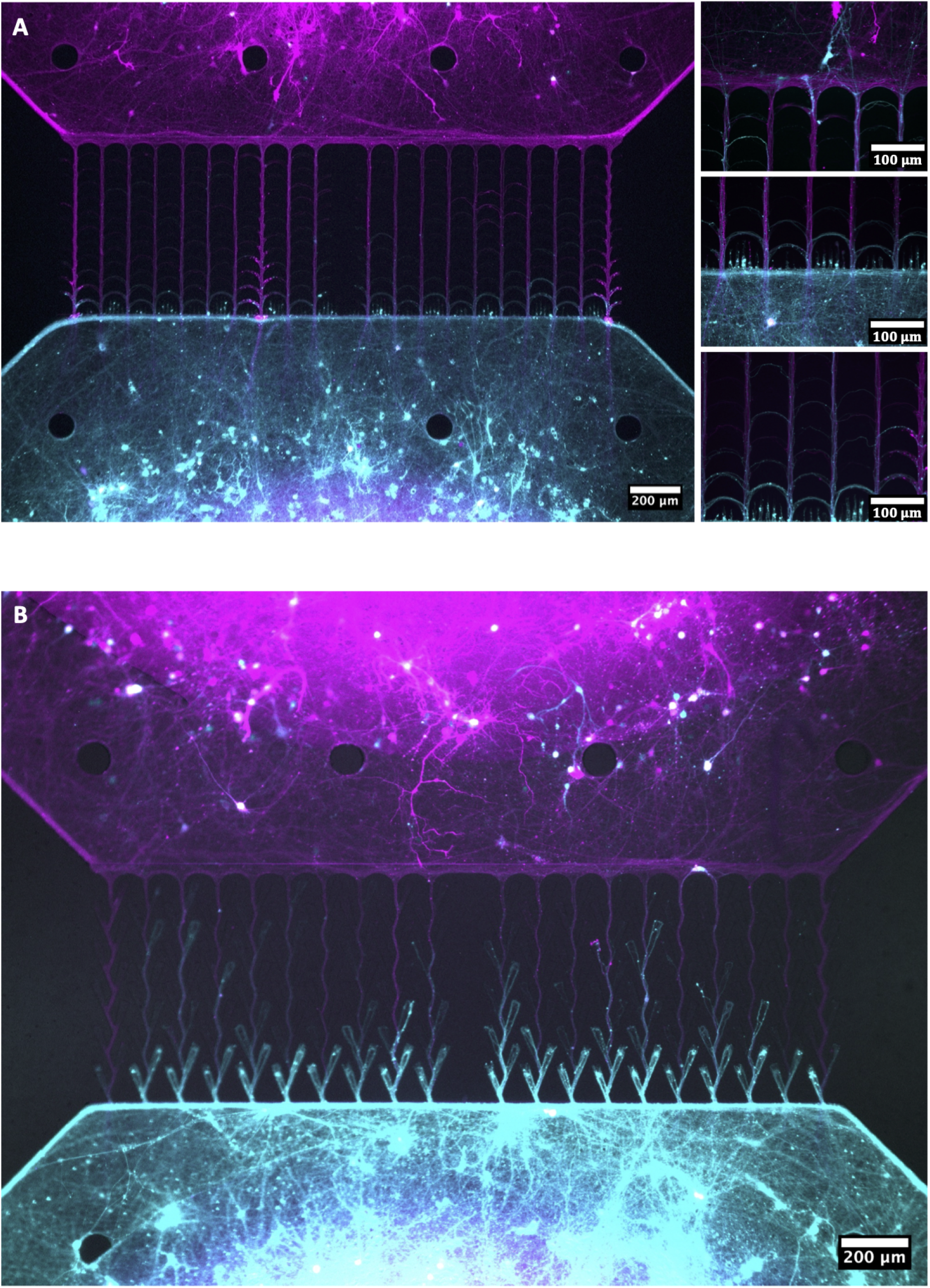
Neuronal cultures transfected with AAV2/1 with a CMV ubiquitous promoter for expression of GFP/mCherry seeded in: **(A.)** U-shapes channels, as pioneered by Renault *et al*. **(B.)** Spine structures as previously described by Gladkov *et al*.

### Testing of Liquid Flow Rate Through Microfluidic Channels

To assess the efficacy of our microfluidic design at keeping the liquid in the two chambers separated, a liquid flow rate test was conducted for both the optimized Tesla valve design (**Figures 3B**, **S2A** & **S3A**) as well as for the bidirectional controls (**Figure S5)**. Both designs had 20 microchannels connecting the two chambers, and each channel was 10 μm wide and 5 μm high. 12 unique microfluidic chips of each design were utilized for the testing.

The flow rate was measured in both the forward (i.e. from presynaptic to postsynaptic chamber) and backward (i.e. from postsynaptic to presynaptic chamber) directions. 75 μL of Neurobasal Plus Medium (Gibco™, A3582801) was added to either the presynaptic or postsynaptic chamber, while the opposing chamber was left empty. After 60 min, a micropipette with 0.5 μL precision was used to measure the amount of liquid that had passed through the channels. Shapiro-Wilk and Shapiro-Francia tests were used to check for normality in the data. As not all data sets were normally distributed, all data is shown using median values, and comparisons between groups were conduced using the Wilcoxon rank sum test.

Box plots with results from the testing are shown in **Figure S7**. Flow rate values were measured to be 1.5 ± 0.71 μL/h and 2.0 ± 0.81 μL/h in the forward direction for the Tesla valve and control microfluidic chips, respectively. In the backward direction, the values were measured to be 1.5 ± 0.66 μL/h and 2.3 ± 0.81 μL/h. Significance (p = 0.027 & p = 0.019 in forward and backward directions, respectively) was found between the Tesla valve microfluidic devices and the controls, but no significance was found between liquid flow in the forward or backward directions for either design.

**Figure S7.**
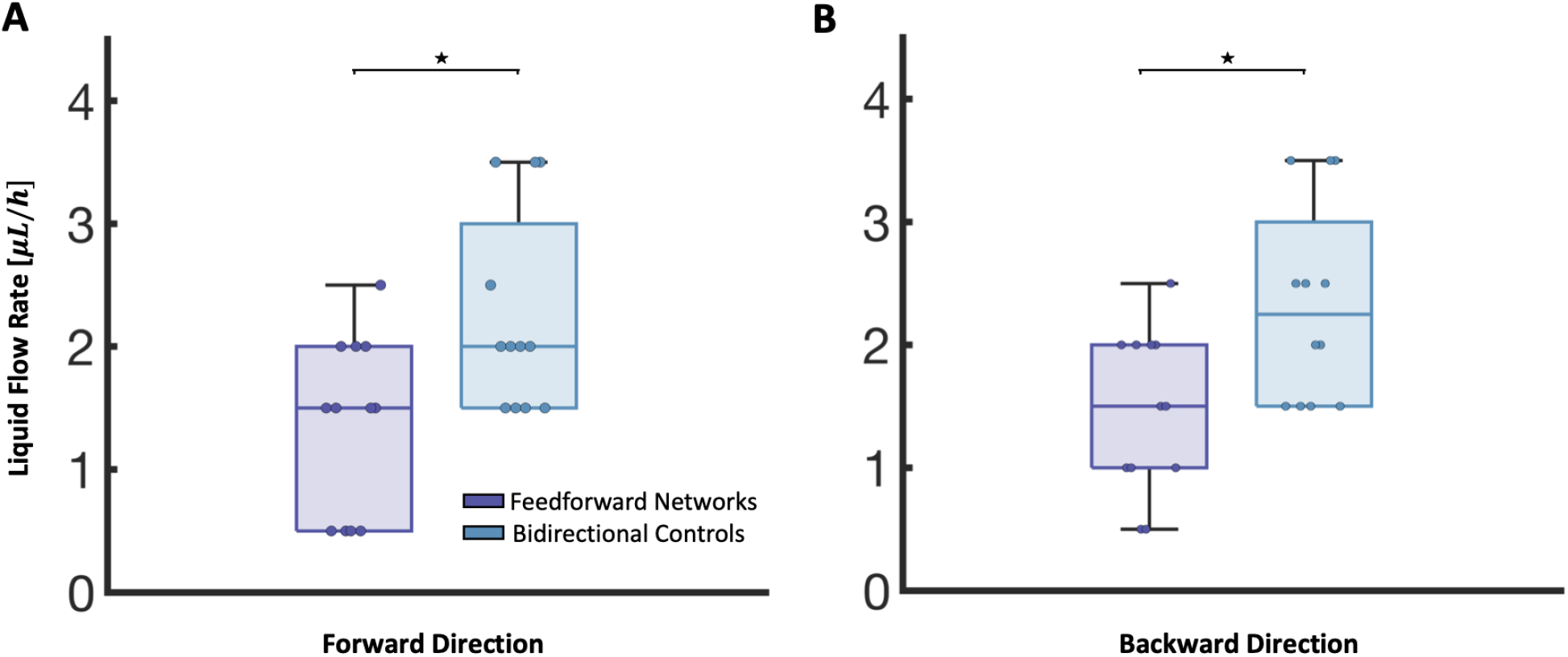
Flow rate testing of microfluidic devices. Box charts showing the rate of liquid flow through the Tesla valve design and bidirectional controls in the forward **(A.)** and backward **(B.)** directions. Significance was assessed using Wilcoxon rank sum test with significance p < 0.05. The charts indicate the median, lower & upper quartiles, and outliers in the data.

It is worth noting that results from this testing are only indicative of the flow rate for the first hour after one chamber is filled up with liquid. As the liquid level in the two chambers equalizes, it would be expected that the flow rate slows down due to the continuous change in the hydrostatic pressure gradient between the chambers. However, the results still show that the liquid flow is minimal between the chambers for both designs. They furthermore indicate that it would take more than 24 h for the liquid level in the two wells to equalize, and that this is irrespective of the flow direction. This is particularly relevant when wanting to keep the conditions in one chamber separated from the other, as during the viral transfections of the cells conducted in this study.

**Figure S8.**
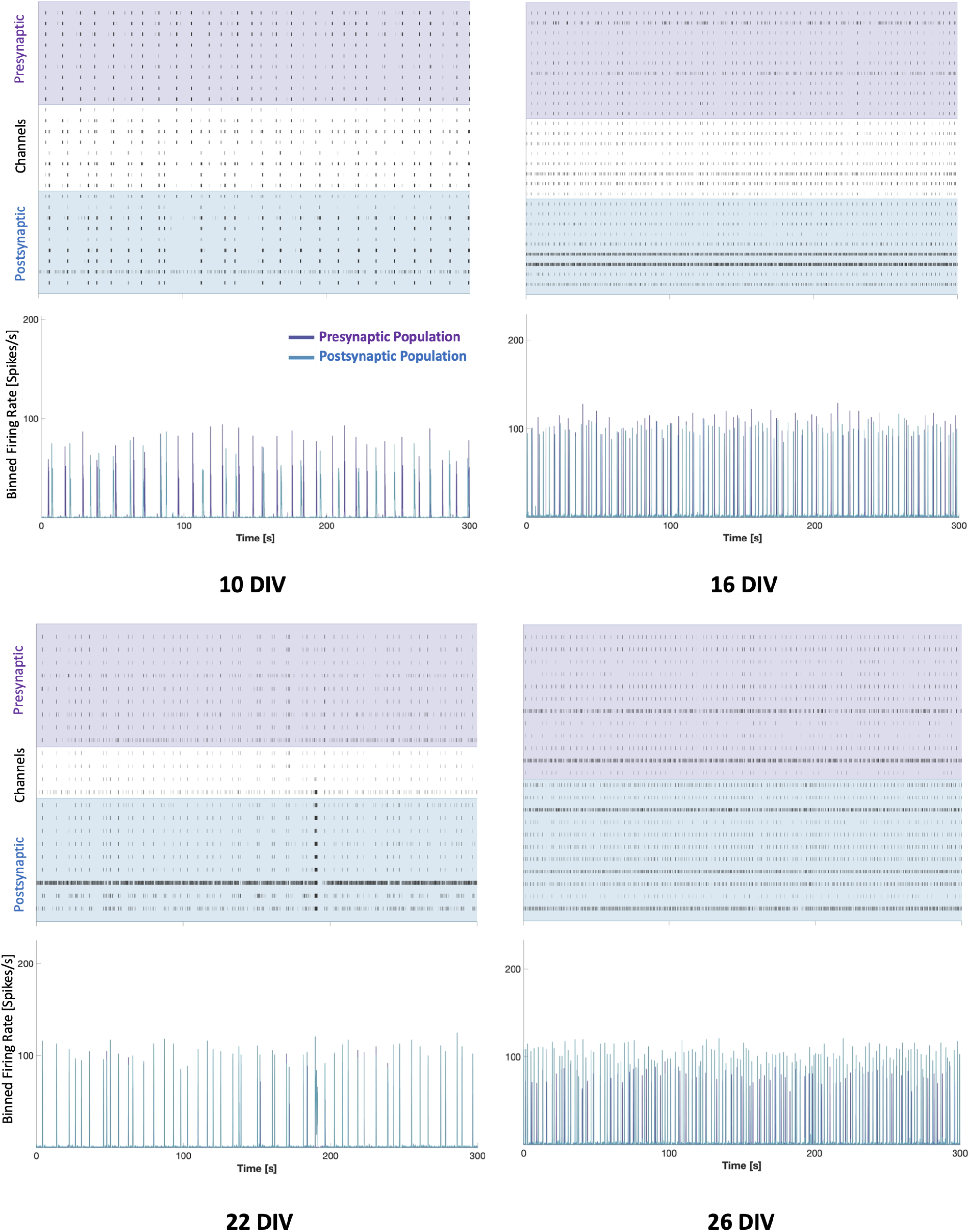
Raster Plots, and 50 ms binned activity plots showing the development in spiking activity for one representative two-nodal network at 10, 16, 22 and 26 DIV, demonstrating the gradual increase in correlated activity between the two nodes before the nodes exhibit more separated activity again at DIV 26.

**Figure S9.**
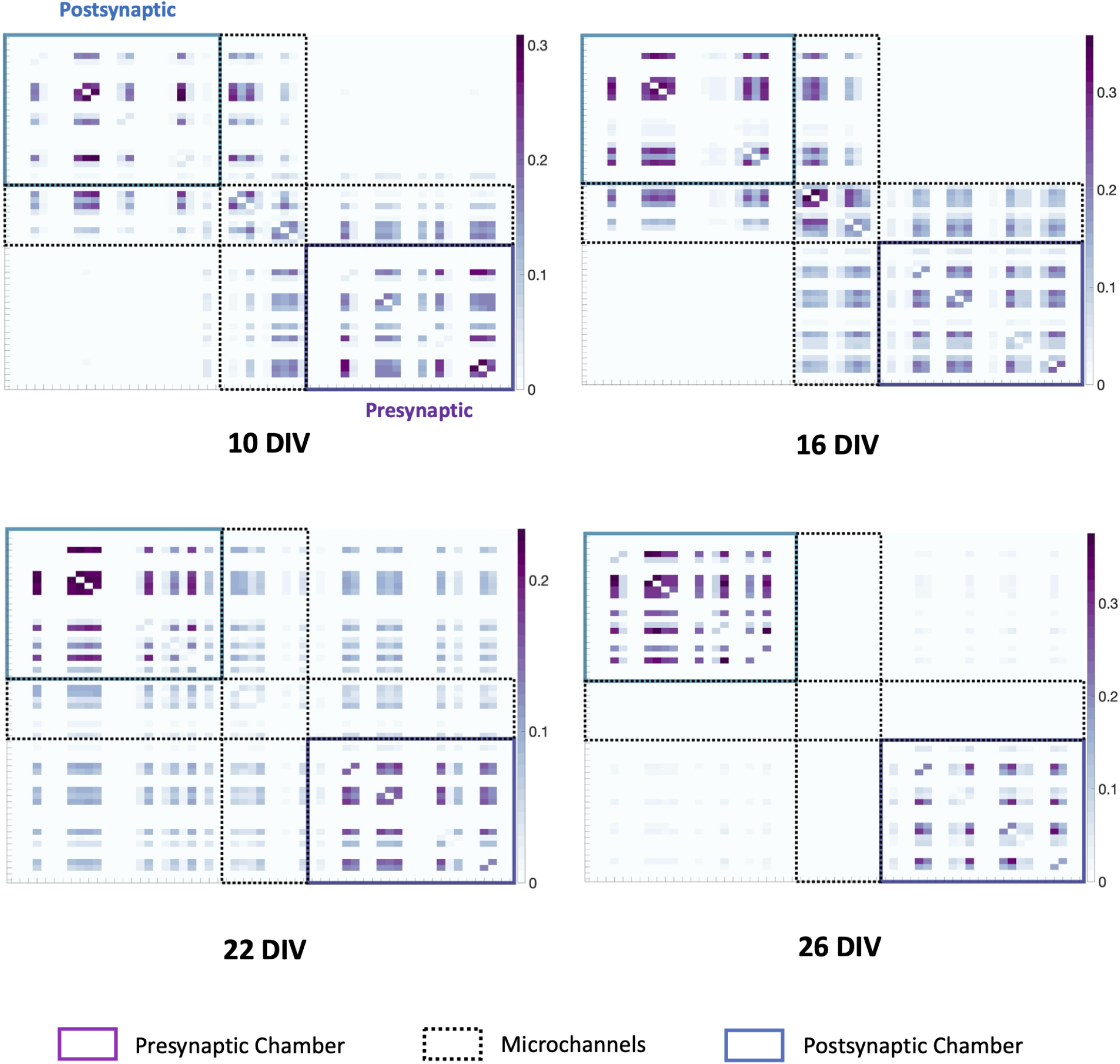
Functional connectivity/adjacency matrices showing the development in functional connectivity for one two-nodal network at 10, 16, 22 and 26 DIV, demonstrating the gradually increasing functional connectivity as the network matures.

**Figure S10.**
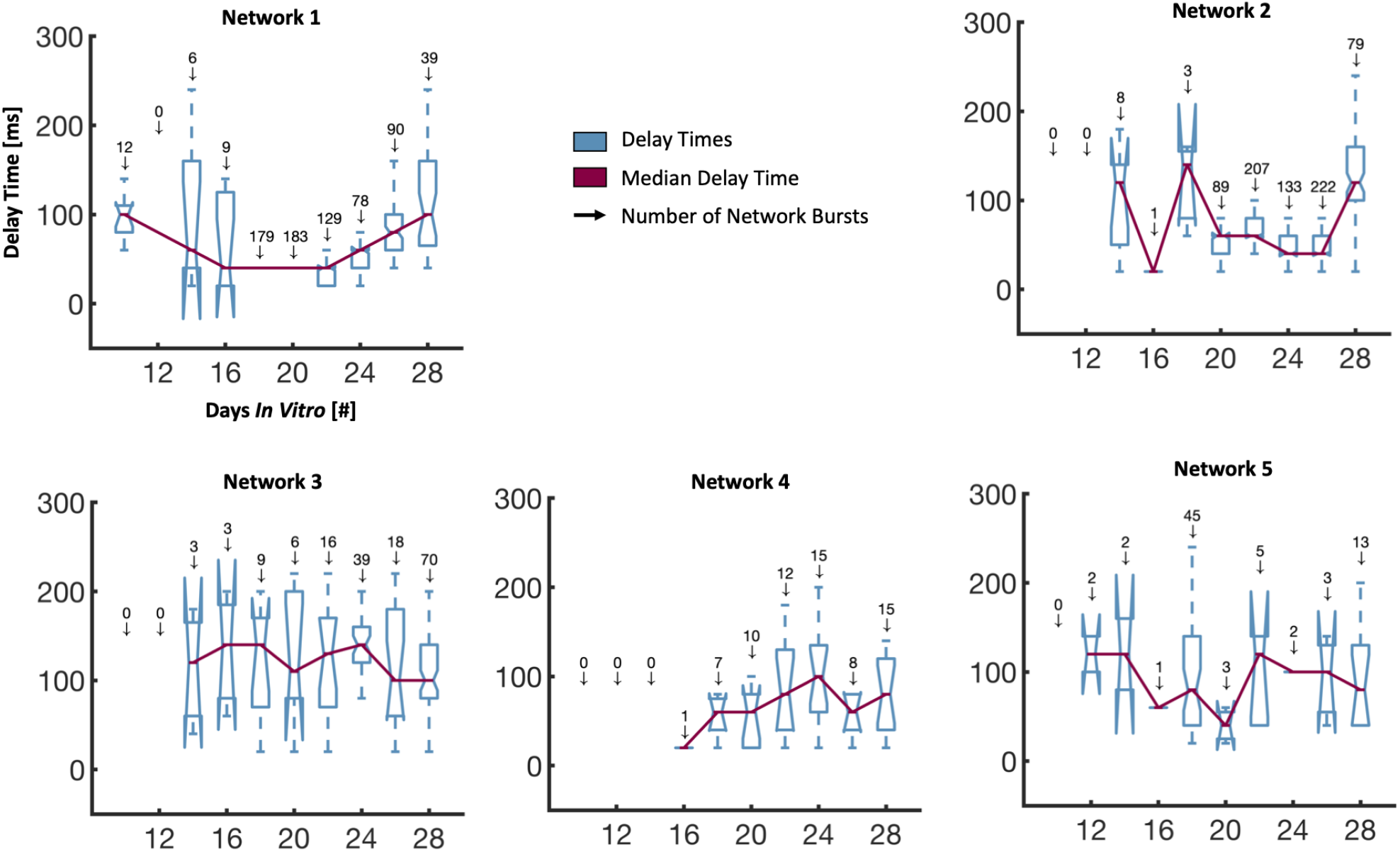
Box charts showing the delay times between the maximum response in the pre- and postsynaptic populations during network bursts for each of the five 2-nodal networks. A full description of the network burst analysis can be found in the main paper. The charts indicate the median, lower & upper quartiles, as well as outliers in the data for each recording day of each network. The number of network bursts detected, and thus analysed, for each recording day is included above the associated box chart.

**Figure S11.**
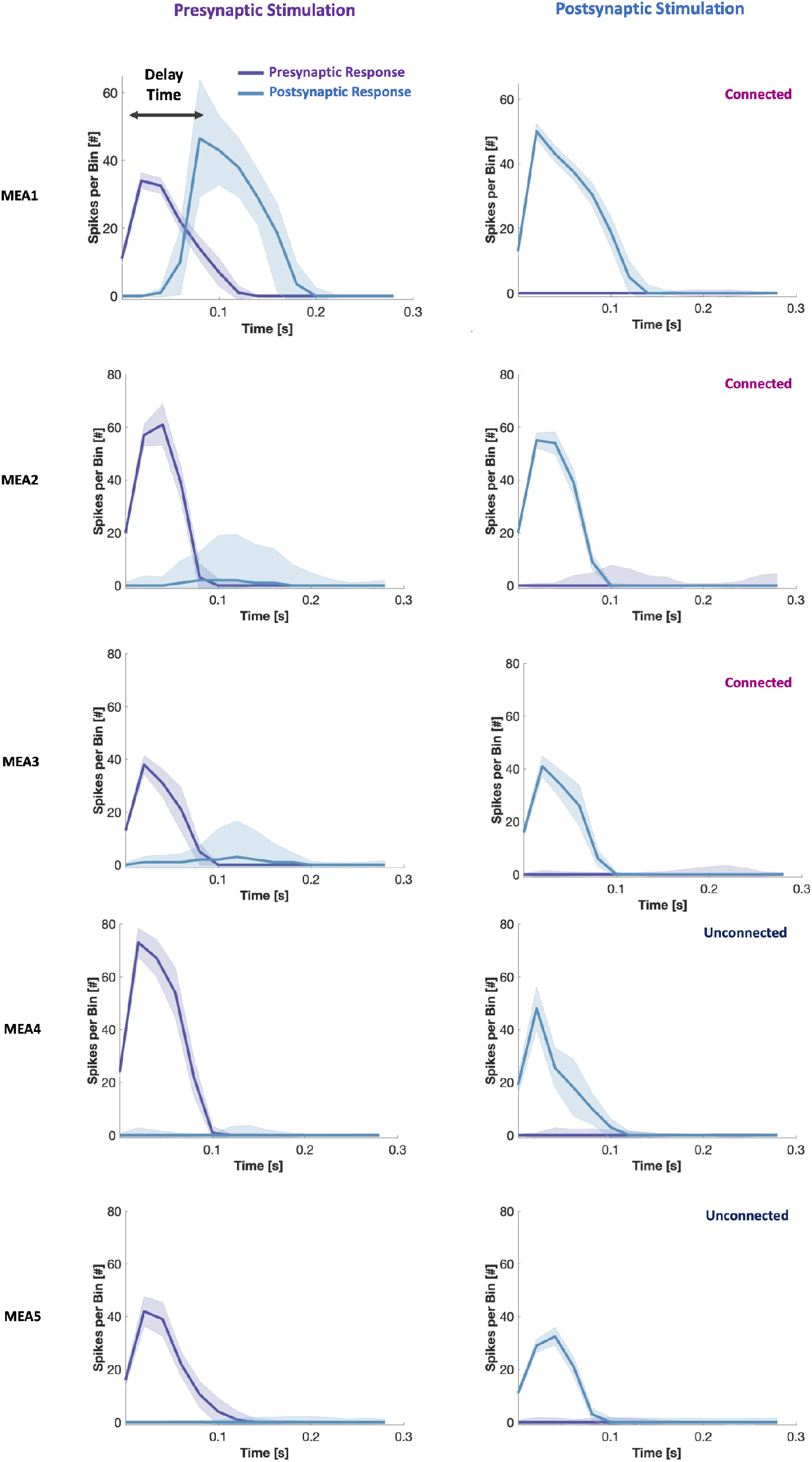
Peristimulus time histograms for the five two-nodal cultures. As can be seen from the graphs, three of the cultures showed a clear spread of activity from the pre-to the postsynaptic population when the presynaptic population was stimulated, while either no or a minimal response was observed in the presynaptic population when the postsynaptic population was stimulated. Two cultures did not appear to have any functional connectivity.

### Protocol for SEM Fixation and Imaging

For cell fixation, all the cell media were removed, and the cultures rinsed with PBS solution two times to remove debris. Consecutively, the PBS was replaced by 2.5% glutaraldehyde in Sorensen’s phosphate buffer (0.1 mol, pH 7.2), barely covering the cells. The cells were incubated in the buffer at 4 °C overnight. Thereafter, a serial dehydration step was conducted to remove all water from the specimens, in steps of 30%, 50%, 70%, 90% and 100% ethanol with 10 min incubation at RT per step. The 100% ethanol was changed 3 times to assure complete removal of all water. Next, a serial drying in hexamethyldisilazane (HMDS, Sigma-Aldrich, 8.04324) was conducted using 33%, 50%, 66% and 100% HMDS, with 5 min for each step. The last step in 100% HMDS was conducted three times to assure removal of all ethanol. Only a thin layer of HMDS barely covering the cells was left in the specimen, and the cultures were left to dry completely out overnight at RT. A 15 nm thick layer of Pt/Pd was sputter coated on the samples prior to imaging (A 208 HR B Sputter Coater, Cressington). The sample was tilted from −45° to 45° with a period of 20 s during the sputtering.

All SEM images were acquired with an APREO Field Emission Scanning Electron Microscope from FEI using the EDT detector to record secondary electrons, with a beam current in the range of 25 – 50 pA, and acceleration voltages between 4.0 – 10.0 k. To avoid build-up of charges, the contact pads of the MEAs were connected to the stage using a copper tape.

